# Molecular Visualization of Neuronal TDP43 Pathology *In Situ*

**DOI:** 10.1101/2024.08.19.608477

**Authors:** Amanda L. Erwin, Martin G. Fernandez, Matthew L. Chang, Durga Attili, Michael Bekier, Emile S. Pinarbasi, Jennifer E. Russ, Renaldo Sutanto, Dafydd Thomas, Xinfei Shen, Ryan D. Baldridge, Elizabeth M. H. Tank, Sami J. Barmada, Shyamal Mosalaganti

## Abstract

Nuclear exclusion and cytoplasmic accumulation of the RNA-binding protein TDP43 are characteristic of amyotrophic lateral sclerosis (ALS) and frontotemporal lobar degeneration (FTLD). Despite this, the origin and ultrastructure of cytosolic TDP43 deposits remain unknown. Accumulating evidence suggests that abnormal RNA homeostasis can drive pathological TDP43 mislocalization, thereby enhancing RNA misprocessing due to the loss of nuclear TDP43, and engendering a cycle that ultimately leads to cell death. Here, we demonstrate that the addition of small monovalent oligonucleotides successfully recapitulates pathological TDP43 mislocalization and aggregation, aberrant splicing, and degeneration in iPSC-derived neurons (iNeurons). By employing a tailored multimodal *in situ* cryo-correlative light and electron microscopy pipeline, we examine the localization and aggregation of TDP43 in near-native conditions. We discover that mislocalized TDP43 accumulates and forms ordered fibrils within autophagosomes and lysosomes in iNeurons, as well as in ALS/FTLD patient tissue. We provide the first high-resolution snapshots of TDP43 aggregates *in situ*, delivering an unprecedented view of the earliest pathogenic events underlying ALS, FTLD, and related TDP43 proteinopathies.

## Introduction

Trans-active response DNA/RNA binding protein of 43 kDa (TDP43) is a ubiquitously expressed nuclear protein responsible for RNA splicing, transport, and stability that is integrally linked with amyotrophic lateral sclerosis (ALS) and frontotemporal lobar degeneration (FTLD)^1–4^. Nearly 95% of individuals with ALS and ∼50% of those with FTLD exhibit nuclear exclusion and cytosolic aggregation of TDP43 within affected neurons and glia^5^. TDP43 mislocalization and cytosolic aggregation are not limited to ALS and FTLD; TDP43 pathology is increasingly recognized as a common event in elderly patients with slowly progressive cognitive decline, termed limbic predominant age-related TDP43 encephalopathy (LATE)^6^. Once excluded from the nucleus, TDP43 primarily accumulates within cytoplasmic aggregates^7,8^. Within cortical regions, these deposits contain amyloidogenic C-terminal TDP43 fragments^9–11^, while spinal cord inclusions are rich in full-length TDP43^12^. These observations suggest that the TDP43 aggregate composition, the mechanisms underlying their formation, and downstream consequences may differ depending on the brain region involved.

Despite its prevalence and increasingly recognized connection with disease pathogenesis, the origin of TDP43 mislocalization and the architecture of native TDP43 inclusions within intact tissue remain mysterious. Engineered mutations disrupting the nuclear localization signal of TDP43^13–15^, application of exogenous stressors^16,17^, addition of recombinant TDP43 fibrils^18,19^, introduction of targeted nuclear export signal nanobodies^20^, or the use of artificial aggregation-promoting domains can all elicit TDP43 mislocalization^21^; yet, these circumstances are disconnected from the pathophysiology of the disease. More recent discoveries suggest that RNA is one of the most significant determinants of TDP43 localization^22–24^. In light of the extensive and widespread abnormalities in RNA processing identified in ALS/FTLD patient material and models^25–27^, this observation raises the intriguing possibility that RNA dysregulation may underlie TDP43 mislocalization and aggregation in these conditions.

Here, we demonstrate that prolonged application of RNA oligonucleotides leads to TDP43 mislocalization, cryptic splicing, neurotoxicity, and the formation of cytosolic TDP43 inclusions in human neurons, resembling aggregates characteristic of human disease. Moreover, cytosolic TDP43 aggregates accumulate primarily within autophagosomes and lysosomes, indicating that these organelles play a crucial role in the formation and/or clearance of mislocalized TDP43. Combining cryogenic-correlative light and electron microscopy (cryo-CLEM) and cryo-electron tomography (cryo-ET), we demonstrate that autophagosomal and lysosomal TDP43 adopts filamentous morphologies mirroring those detected in ALS and FTLD with TDP43 pathology (FTLD-TDP)^28,29^. In keeping with these observations, we observe abundant TDP43 filaments within ALS/FTLD patient tissue-derived lysosomes. Based on these findings, we propose that RNA dyshomeostasis, in conjunction with ineffective autophagy or lysosomal turnover — processes often impaired in aging^30–37^ — could contribute to nuclear TDP43 mislocalization and the accumulation of fibrillar aggregates characteristic of TDP43 proteinopathies such as ALS and FTLD-TDP.

## Results

### Establishing an in vitro neuronal model to recapitulate TDP43 pathology

Short oligonucleotides (oligos) bearing GU motifs, which are preferentially recognized by TDP43, effectively promote TDP43 nuclear efflux in transformed cell lines^22–24^. To study a cell type relevant to ALS and FTLD, we applied short, monovalent GU-rich RNA (5’-GUGUGUGUGUGU-3’, (GU)_6_ or (GU)_6_ oligos, hereafter) to human induced pluripotent stem cell (iPSC) derived neurons (iNeurons). All oligonucleotides were modified by 2’-*O*-methyl and phosphorothioate groups to enhance resistance to RNase degradation. A customized cassette encoding the master transcription factors NGN1 and NGN2, under the control of a doxycycline (dox)-inducible promoter, was integrated into the *CLYBL* safe harbor locus of iPSCs, facilitating robust and rapid differentiation of iPSCs into glutamatergic, forebrain-like iNeurons (**Figure 1A**)^26,38–43^. (GU)_6_ oligos were transfected using Endo-Porter before assessing TDP43 localization by immunostaining and fluorescence microscopy. In contrast to the rapid (<6h) effect of RNA oligos on TDP43 localization in transformed cell lines^22–24^, we observed pronounced mislocalization and cytosolic accumulation of TDP43 in iNeurons only after 24h or more of (GU)_6_ treatment (**Figures 1B and 1C; Figures S1C and S1F**). We did not observe such mislocalization in untreated iNeurons or those exposed to Endo-Porter alone or Endo-Porter with control RNA oligos (5’-CCCCCCCCCCCC-3’, C_12_ hereafter) (**Figures 1B-1D and S1A-S1F**). By 24h post-(GU)_6_ treatment, approximately half of all iNeurons displayed a greater than 50% reduction in the nuclear-to-cytoplasmic (N/C) ratio of TDP43 (**Figures 1C and 1D**). Based on these kinetics, we chose the 24h time point for all subsequent experiments.

**Figure 1.**
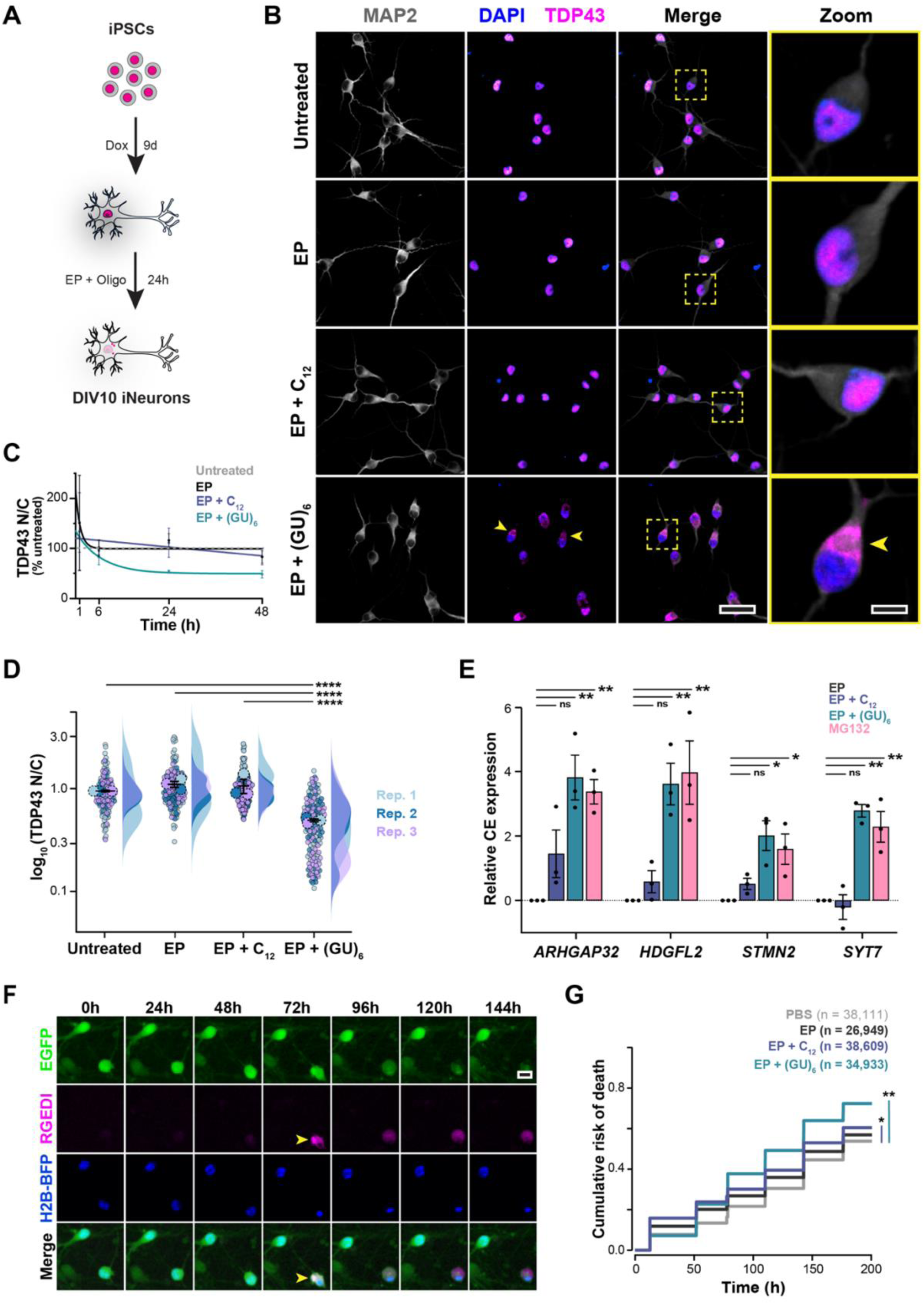
Short GU-rich oligonucleotides induce TDP43 nuclear export, cryptic exon splicing, and neurotoxicity in human iNeurons. **(A)** Schematic outline of RNA-induced TDP43 nuclear egress and cytoplasmic deposition in iPSC-derived neurons. **(B)** Representative confocal fluorescence microscopy images of untreated DIV10 iNeurons (top row) or iNeurons treated with Endo-Porter only (EP, second row), Endo-Porter with control oligo (EP + C_12_, third row), or Endo-Porter with GU-oligo (EP + (GU)_6_, last row) for 24h before being chemically fixed and stained for neuronal marker (MAP2), nuclei marker (DAPI), and TDP43. TDP43 mislocalization from the nucleus to cytoplasm was evident only in (GU)_6_-treated intact iNeurons (last row); yellow arrowheads highlight select iNeurons. A representative iNeuron (yellow dashed box) for each condition is shown at 5× magnification (last column). Images are pseudo-colored as follows: MAP2 (gray), DAPI (blue), and TDP43 (magenta). Scale bar: 50µm, except for 5× magnified image, which is 10µm. **(C)** TDP43 nucleus vs. cytoplasm (N/C) fluorescence intensity ratios in DIV10 iNeurons either left untreated (gray dashed line) or treated with EP alone (black), EP with 2µM C_12_ (purple), or EP with 2µM (GU)_6_ (teal) for 1-48h. N/C ratios for each treatment and time point were normalized to the mean value of the untreated control and reported as a percentage of the untreated control. (mean 土 SEM, n > 140 across three biological replicates for each treatment and time point, curves were fit to the mean values using non-linear regression with a one-phase exponential decay model). **(D)** Log-transformed ratio of TDP43 fluorescence intensity in the nucleus vs. cytoplasm (N/C ratio) plotted for DIV10 iNeurons either untreated or treated with EP only, EP with 2µM C_12_, or EP with 2µM (GU)_6_ for 24h. TDP43 N/C ratios were normalized to the untreated condition for each replicate. (mean 土 SEM, Kruskal-Wallis test followed by Dunn’s post hoc test, ****p < 0.0001, n > 148 across three biological replicates). Biological replicates 1-3 are colored as light blue, blue, and purple, respectively. **(E)** Quantification of the de novo cryptic exon inclusions, by RT-qPCR analysis for a panel of TDP43 substrates, *ARHGAP32, HDGFL2, STMN2, SYT7* in DIV10 iNeurons treated with either EP (black), EP and 2 µM C_12_ (EP + C_12_, purple), EP and 2 µM (GU)_6_, (EP + (GU)_6_, teal), or 100nM of potent proteasome inhibitor, MG132 (pink), for 24h. Transcript expression levels were normalized to *GAPDH* transcript levels and the Endo-Porter treated condition for each replicate (mean 土 SEM, Tukey test, **p < 0.01, *p < 0.05, n = 3). **(F)** Human iNeurons were tracked daily for 10d by automated microscopy, visualized using a cell marker (EGFP) in green, nuclear marker (H2B-BFP) in blue, and a red genetically encoded death indicator (RGEDI) in red. Yellow arrowheads at 72h indicate time of death, marked by increases in RGEDI fluorescence, rounding of the cell body, and retraction of neurites. **(G)** Cumulative risk of death in human iNeurons treated with PBS (gray), EP alone (black), or EP + C_12_ (purple) or (GU)_6_ (teal) oligos (2µM each). n = number of neurons, combined from 6 technical replicates per each of the 3 biological replicates. * Hazard ratio (HR) = 1.09, p <2×10^-16^; ** HR= 1.22, p <2×10^-16^; Cox proportional hazards analysis.

Loss of nuclear TDP43 in disease results in distinct splicing changes, including the abnormal inclusion of unannotated (cryptic) exons in TDP43 target RNAs. We therefore assessed whether treatment with (GU)_6_ oligos triggered cryptic splicing of canonical TDP43 substrates affected in ALS and FTLD-TDP^44,45^. Cryptic products of *ARHGAP32, HDGFL2, STMN2,* and *SYT7* were dramatically upregulated in iNeurons upon treatment with (GU)_6_ for 24h (**Figure 1E**), consistent with a decrease in the TDP43 nuclear-to-cytoplasmic ratio in these cells (**Figures 1C and 1D**). We also observed significant cryptic splicing with the proteasome inhibitor MG132, in agreement with the rapid mislocalization of TDP43 triggered by this compound^46,47^. These results confirm that RNA-induced TDP43 mislocalization in iNeurons is associated with the loss of nuclear TDP43 splicing activity.

A central and defining feature of neurodegenerative diseases such as ALS and FTLD is the progressive loss of neurons in affected brain regions. To determine whether (GU)_6_-dependent TDP43 mislocalization leads to neuron loss, we employed automated longitudinal microscopy to assess the fate of individual iNeurons treated with oligos^26,48^. iNeurons treated with Endo-Porter alone, Endo-Porter + C_12_, or Endo-Porter + (GU)_6_ oligos were imaged at regular 24h intervals over 10 days by automated microscopy. We applied semantic image segmentation algorithms to detect individual neurons, and defined the time of death using a red-shifted genetically encoded death indicator (RGEDI)^49^ that fluoresces only when intracellular Ca^2+^ levels rise rapidly. Cell death was also characterized by morphological changes, such as rounding of the soma, neurite retraction, and blebbing (**Figure 1F**). Using this assay, we found that treatment with either C_12_ and (GU)_6_ oligos significantly increased the risk of death compared to no treatment or Endo-Porter alone, suggesting mild non-specific toxicity associated with RNA transfection in neurons (**Figure 1G**). Consistent with the selective impact of (GU)_6_ oligos on TDP43 localization, however, we observed the highest risk of death in neurons treated with (GU)_6_ (**Figure S1G**).

We also examined oligo-treated iNeurons for post-translational modifications typical of mislocalized TDP43 in ALS and FTLD-TDP. Consistent with pathological TDP43 aggregates^3,7,8,50^, cytoplasmic TDP43 puncta were phosphorylated at S409/S410. We observed a pronounced increase in phosphorylated TDP43 (pTDP43) levels in (GU)_6_-treated iNeurons after 10 days, suggesting that phosphorylation is a delayed event rather than a primary driver of TDP43 aggregation (**Figures S2A-C**). Chronic (GU)_6_ exposure for 10 days led to a further decrease in the TDP43 N/C ratio, reflecting the accumulation of large cytoplasmic aggregates positive for both TDP43 and pTDP43 (**Figures S2A, S2B, and S2D**). Taken together, these findings indicate that RNA-induced perturbation of TDP43 localization in iNeurons recapitulates key features of ALS/FTLD-TDP pathology, including TDP43 nuclear egress, cryptic exon inclusion, TDP43 phosphorylation, and neuronal loss^7,44,45,51^.

### A custom cryo-CLEM workflow to visualize mislocalized TDP43 in human neurons

Given the strong genetic and mechanistic links between ALS/FTLD and dysfunction of the autophagosome–lysosome system^52–55^, we hypothesized that these organelles may play a pivotal role in the sequestration and clearance of mislocalized TDP43. To test this, we assessed the colocalization of TDP43 and the lysosomal marker LAMP1 in iNeurons treated for 24h with either C_12_ or (GU)_6_ oligos using confocal microscopy **(Figure S3)**. Consistent with previous observations, (GU)_6_-treated iNeurons displayed a significant decrease in the TDP43 N/C ratio (**Figures S3A and S3B**). Total LAMP1 levels, as measured by mean fluorescence intensity, were comparable between C_12_ and (GU)_6_-treated iNeurons (**Figure S3C**). Notably, (GU)_6_ treatment led to a significant increase in overlap between TDP43 and LAMP1, suggesting lysosomal accumulation of mislocalized TDP43 (**Figures S3D**).

While light microscopy enables tracking of protein and organelle dynamics, it lacks the ultrastructural resolution required to visualize the subcellular context and organization of pathological protein assemblies. For example, light microscopy alone cannot reveal the molecular characteristics of lysosomal and peri-lysosomal deposits (**Figure S3A**, white and orange arrowheads, respectively). To overcome this limitation and interrogate the native architecture of mislocalized TDP43 in human iNeurons, we established a robust *in situ* cryo-correlative light and electron microscopy (cryo-CLEM) workflow. This approach integrates the molecular specificity of fluorescence imaging with the nanometer resolution of cryo-electron tomography (cryo-ET), enabling precise spatial correlation of TDP43 within an intact cellular environment (**Figure 2**). To specifically visualize neuronal lysosomes with cryo-CLEM, iPSCs were differentiated into iNeurons as before (**Figure 1A**), but on micropatterned gold electron microscopy grids (**Figure 2A**, schematic to the left). The grids are manually screened for consistency in neuron distribution and quality (cell density, presence of neurites, etc.) using a standard widefield microscope. This is crucial for imaging and achieving high-resolution insights from morphologically healthy neurons, as grids containing iNeurons, unlike those with other mammalian cell types, exhibit a high degree of heterogeneity (**Figure 2A**, right). After treating iNeurons with Endo-Porter and (GU)_6_ for 24h, we added LysoTracker Green DND-26, a fluorescent weak base that selectively accumulates in acidic organelles such as lysosomes. Due to the properties of the dye and the dynamic behavior of the endolysosomal system, LysoTracker also labels other acidified organelles in this pathway. As these structures are morphologically indistinguishable in cryo-electron tomograms, we will collectively refer to endolysosomes, autolysosomes, and terminal lysosomes as lysosomes throughout this study. Within 5m of the addition of LysoTracker, FluoSpheres were added for fiducial-based correlation, and the grids were plunge-frozen in liquid ethane. LysoTracker was employed to visualize lysosomes with cryo-CLEM, taking advantage of the fact that dyes are far brighter and yield less background than GFP or its spectral variants. We collected cryo-confocal z-stacks of candidate iNeurons (**Figure 2B**) and transferred autogrids containing cells to a dual-beam cryo-focused ion beam scanning electron microscope (cryo-FIB/SEM). We then used the 3D Correlation Toolbox (3DCT) to precisely target the region of interest based on the cryo-confocal images (**Figures 2B and 2C**)^56^. Following cryo-focused ion beam (cryo-FIB) milling, the lamellae were assessed by cryo-transmission electron microscopy (cryo-TEM), allowing for a second correlation of the fluorescence plane, previously identified from 3DCT correlation (**Figure 2C**), with the medium magnification cryo-TEM image (6500×, **Figure 2D**). Tilt series data collection, followed by tomogram reconstruction on areas guided by the fluorescence signal, revealed the presence of filaments within lysosomes of (GU)_6_-treated iNeurons (**Figure 2E and Video S1)**.

**Figure 2.**
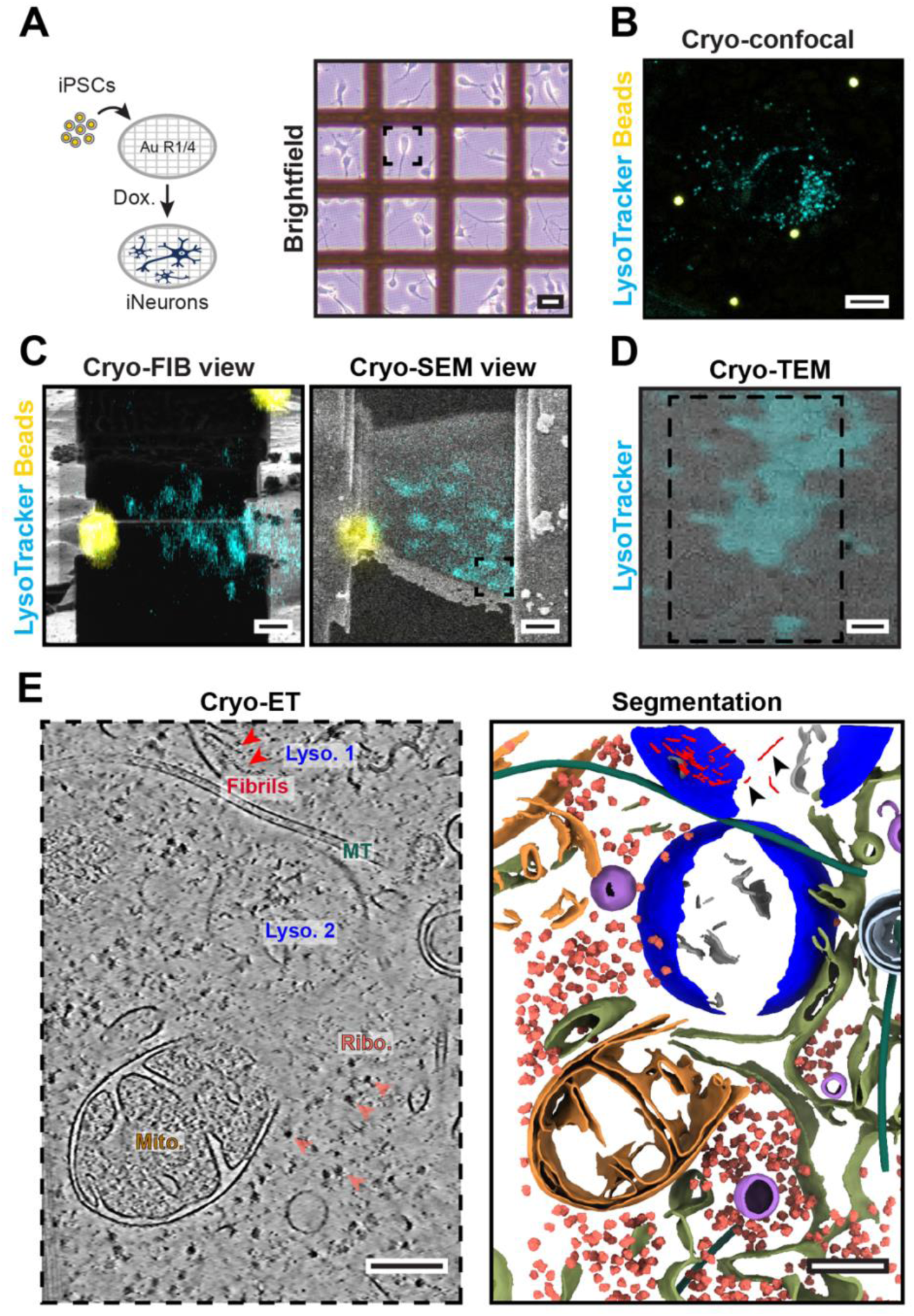
*In situ* cryo-CLEM workflow for ultrastructural analysis of human iNeurons. (**A**) Schematic of doxycycline-inducible iPSCs differentiated to iNeurons on micropatterned electron microscopy grids (left). The grids are manually checked under a widefield microscope for quality and are subsequently plunge-frozen in liquid ethane. Representative widefield image of iNeurons growing on micropatterned grids (right). Scale bar: 50µm. Endo-Porter and (GU)_6_ oligos are added to the grids 24h before plunge-freezing, LysoTracker is added 5m before, and FluoSpheres (diameter - 2µm) are added immediately before plunge-freezing. (**B**) A pseudocolored maximum intensity projection image of a cryo-confocal z-stack with LysoTracker (cyan), and FluoSphere beads (yellow) collected from an iNeuron (black dashed box in panel A) on a grid for precise correlative cryo-FIB/SEM milling. Scale bar: 10µm. (**C**) Overlay of the cryo-confocal fluorescence microscopy data onto a lamella milled by correlation through the 3D-correlation toolbox (3DCT). The cryo-FIB view (left) and cryo-SEM view (right) of the same lamella are shown. Scale bar: 2µm. (**D**) Overlay of the z-slice of LysoTracker fluorescence (cyan) and 6500× cryo-TEM montage image of the lamella acquired at the region highlighted in the cryo-SEM view image (black dashed box in panel C). Tomography tilt series are collected in the areas of the cryo-lamella highlighted (black dashed rectangle). Scale bar: 200nm. (**E**) Slice through a reconstructed and denoised tomogram showing intact neuronal interiors (left). Neuronal microtubule (MT), lysosomes (Lyso. 1 & 2), mitochondria (Mito.), and ribosomes (Ribo.) are readily visible. The fibrils within the top lysosome and the ribosomes are highlighted with red and salmon arrowheads, respectively. Scale bar: 200nm. The corresponding 3D segmentation (right) highlights cellular features rendered as: ribosomes (salmon), fibrils (red), endoplasmic reticulum (olive green), mitochondria (golden orange), microtubules (dark green), lysosome (blue), membrane cargo (gray), vesicles (lilac), and membrane whorls/multivesicular bodies/intralumenal vesicles (light blue). Scale bars: 200nm. Fibrillar deposits (black arrowheads) are only observed in one of the two lysosomes in the field of view (Lyso. 1, top).

### In situ TDP43 architecture at molecular resolution

Lysosomes are dynamic, heterogeneous organelles whose contents may vary considerably even within the same compartment and subcellular location^57^. Evidence for this is clear in **Figure 2 and Video S1**; although the field of view contains two lysosomes marked by LysoTracker fluorescence (**Figure 2D**), several fibrils are visible in the lysosome at the top of the field, while the centrally located lysosome is devoid of any fibrillar material (**Figure 2E**, *Lyso. 1 vs. Lyso. 2*). To determine if these filaments are comprised of TDP43, we employed CRISPR/Cas9 to insert an open reading frame encoding HaloTag immediately downstream of the endogenous *TARDBP* start codon in human iPSCs, thereby generating a fusion of HaloTag with the N-terminus of native TDP43 (**Figure S4A**). The correct positioning of HaloTag was confirmed by Sanger sequencing, and an appropriate shift in the molecular weight of TDP43 was observed by immunoblotting (**Figure S4B**). After integrating the *NGN1/NGN2* cassette, HaloTag-TDP43 iPSCs were differentiated into iNeurons as above and labeled using a cell-permeable and photostable HaloTag-compatible dye (JF635)^58^. We then applied GU_6_ oligos for 24h and employed our cryo-CLEM workflow to visualize mislocalized, JF635-positive HaloTag-TDP43 puncta within the cytoplasm in greater detail (**Figures S4C-4E**). Many of these HaloTag-TDP43 foci formed accumulations of various sizes that excluded ribosomes and other cellular organelles, consistent with the characteristic nature of phase-separated condensates. (**Figure 3A and Video S2**). Notably, such ‘ribosome-excluded zones’ (hereafter, exclusion zones or condensates) are frequently localized to the surface of lysosomes, where they adopt the curvature of the adjacent membrane (**Figure 3A**, cyan and yellow arrowheads). HaloTag-TDP43 positive lysosomes near these exclusion zones contained fibrils (**Figure 3A**, red arrowheads). Among these exclusion zones, some appeared to be ‘amorphous’ with very little organized ultrastructure (**Figure 3A**, yellow boundaries), and some of these exclusion zones exhibited different material properties and had a visible semi-ordered arrangement with interspersed smaller filaments, the exact composition of which remains to be determined (**Figure 3A**, cyan boundary and **Video S2**). These latter regions are remarkably similar to solid-like protein-RNA condensates noted in previous studies^59^.

**Figure 3.**
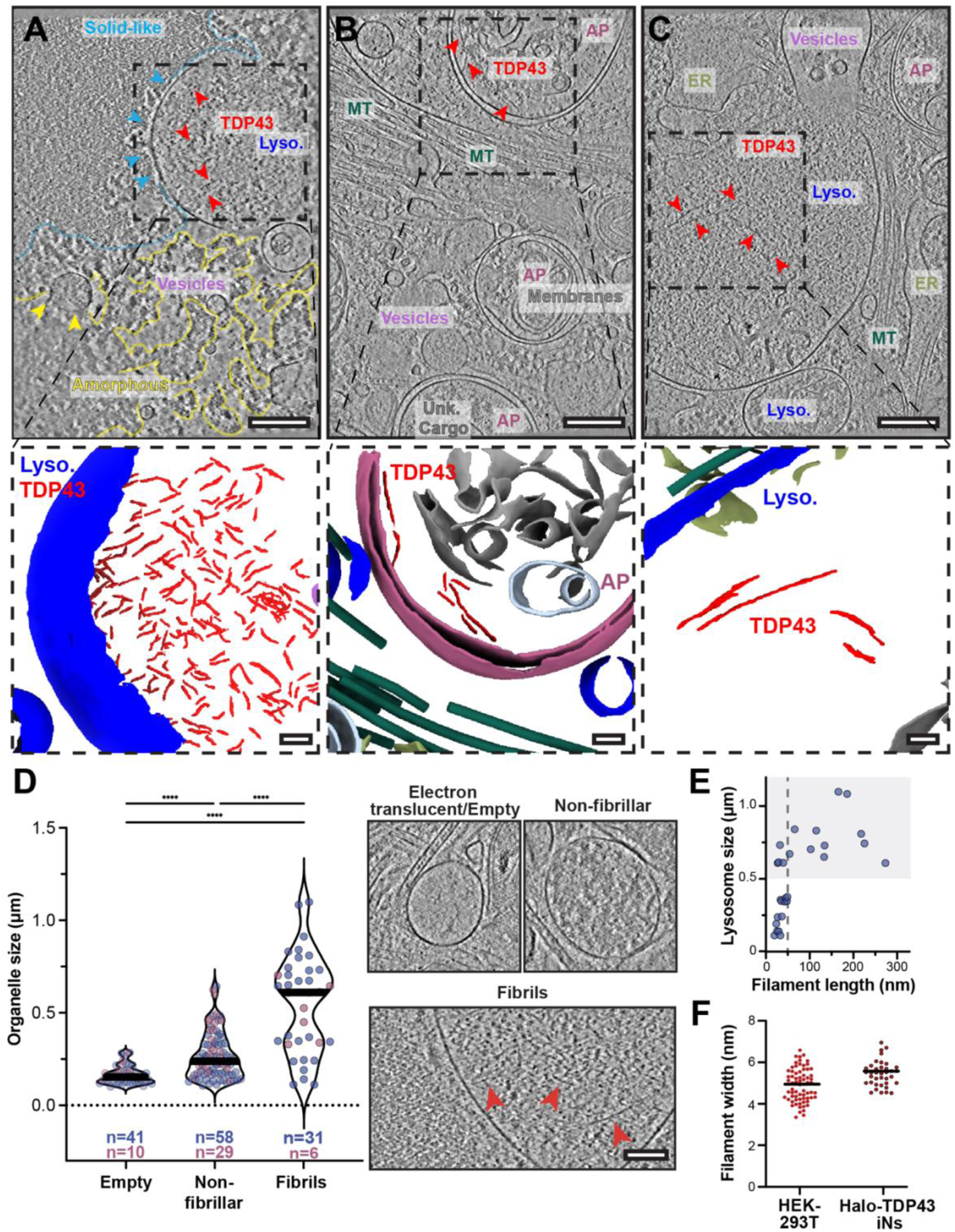
Mislocalized TDP43 accumulates within neuronal autophagosomes and lysosomes. (**A-C**) Top: Slice through reconstructed tomograms from HaloTag-TDP43 iNeurons treated with EP and (GU)_6_ oligo showing intact neuronal interiors. Neuronal lysosomes (Lyso.), vesicles, microtubules (MT), autophagosomes (AP), endoplasmic reticulum (ER), condensate exclusion zones (amorphous or solid-like), membranes, and TDP43 (red arrowheads) are highlighted. A representative example of a condensate proximal to a lysosome containing shorter TDP43 fibrils is shown in (**A**), autophagosomes that have engulfed TDP43 fibrils in (**B**), and an enlarged lysosome with TDP43 fibrils is shown in (**C**). Scale bars: 200nm. Bottom: The corresponding segmentation of the tomograms showing the zoomed-in view of the sub-region highlighted by the dashed black box. Cellular features rendered as: TDP43 (red), endoplasmic reticulum (olive green), microtubules (dark green), lysosome (blue), membranes (gray), multivesicular bodies, membrane whorls or intralumenal vesicles (light blue), and autophagosomes (purple) are highlighted. Scale bars: 100 nm. (**D**) Left: Quantification of organelle size with respect to the luminal content. Single-membraned lysosomes, either larger than 100nm (indicated by blue circles), and double-membrane bound autophagosomes (indicated by purple circles) were categorized from ∼50 tomograms reconstructed from tilt series acquired on lamellae from HaloTag-TDP43 iNeurons treated with EP and (GU)_6_ oligos. These organelles were classified based on their internal luminal contents as devoid of electron-dense material within the lumen (electron translucent), containing unknown cargo, amorphous or membranous material (non-fibrillar), and bona fide fibrils (fibrils). Medians are shown as thick lines. Right: Representative slices through tomograms of electron translucent, non-fibrillar, or fibril-containing single-membrane-bound organelles. (**E**) Quantification of lysosome size with respect to TDP43 filament length (n = 28). The gray dashed line indicates 50nm. The gray box encompasses lysosomes larger than 0.5µm. (**F**) Distribution of lysosome-bound filament widths from lysosomes isolated from HEK293T cells (mean 土 SD = 4.9 土 0.79nm, n = 69) and lysosomes within Halo-TDP43 iNeurons (mean 土 SD = 5.5 ± 0.61nm, n = 37).

Furthermore, we observed cytosolic TDP43 foci located within double membrane-bound structures reminiscent of autophagosomes (**Figure 3B and Video S3**). TDP43-containing autophagosomes also exhibit a variety of other cellular debris or amorphous material, consistent with bulk (macro) autophagy of TDP43.

A significant portion of mislocalized TDP43 appeared to be fibrillar rather than amorphous and was present within single, membrane-bound lysosomes (**Figure 3C and Video S4**). Within these organelles, the density and size of TDP43 filaments varied considerably (**Figure 3C**). Overall, we observed TDP43 fibrils within ∼25% of imaged lysosomes (n = 133) and ∼12% of autophagosomes (n = 46). We classified the autophagosomes and lysosomes based on the luminal content into either ‘empty’, electron translucent with no visible electron dense material within the lumen; ‘non-fibrillar’, containing amorphous or membranous material; or ‘fibrillar’, containing bona fide filaments or fibrils (**Figures S5A and S5B**). We did not find long, ordered TDP43 filaments outside of acidified membrane-enclosed organelles such as lysosomes and autophagosomes, suggesting that fibril formation may be enhanced by low pH, molecular crowding, and/or partial cleavage of TDP43 within these organelles^60^.

We next assessed the size of fibril-containing lysosomes across all cryo-tomograms collected of Halo-TDP43 iNeurons treated with (GU)_6_ oligos. Despite the broad distribution of lysosome size, fibril-containing lysosomes appeared significantly larger than empty or non-fibrillar lysosomes (**Figure 3D**). We observed a clear correlation between fibril length and the size of autophagosomes and lysosomes (**Figure 3E**). The average fibril width of 5.5 ± 0.61nm (**Figure 3F**) closely matches the mean width of amyloidogenic TDP43 C-terminal fibrils obtained by proteinase K-mediated digestion (5.4 ± 1.5nm), as well as that of pronase-treated sarkosyl insoluble fibrils obtained from FTLD Type A, suggesting that intralysosomal TDP43 fibrils undergo partial proteolysis^28,61^.

### Mislocalized TDP43 accumulates within autophagosomes and lysosomes in other cell types

TDP43 mislocalization is not limited to neurons in ALS and FTLD-TDP but is also found in glia and other CNS cell types^5^. We therefore investigated whether TDP43 demonstrates similar accumulation within autophagosomes and lysosomes of non-neuronal cells upon treatment with (GU)_6_ oligos. To achieve this, we utilized an engineered HEK293T cell line in which the endogenous *MAP1LC3B* gene was fused to an open reading frame encoding the photoconvertible fluorescent protein Dendra2^40^. As in iNeurons, we transfected Dendra2-LC3B HEK293T cells with Endo-Porter alone or Endo-Porter + (GU)_6_ oligo. After 4h, we isolated Dendra2-positive autophagosome-rich and lysosomal-rich fractions by density gradient centrifugation and immunoblotted for TDP43 content within each fraction (**Figures 4A and S6A**). Cells transfected with Endo-Porter + (GU)_6_ oligos displayed a significant shift in the distribution of TDP43 towards lysosomal and autophagosomal fractions (**Figure 4A**) that was not observed in HEK293T cells transfected with Endo-Porter alone (**Figure 4A**), consistent with RNA-induced accumulation of TDP43 within lysosomes and autophagosomes.

**Figure 4.**
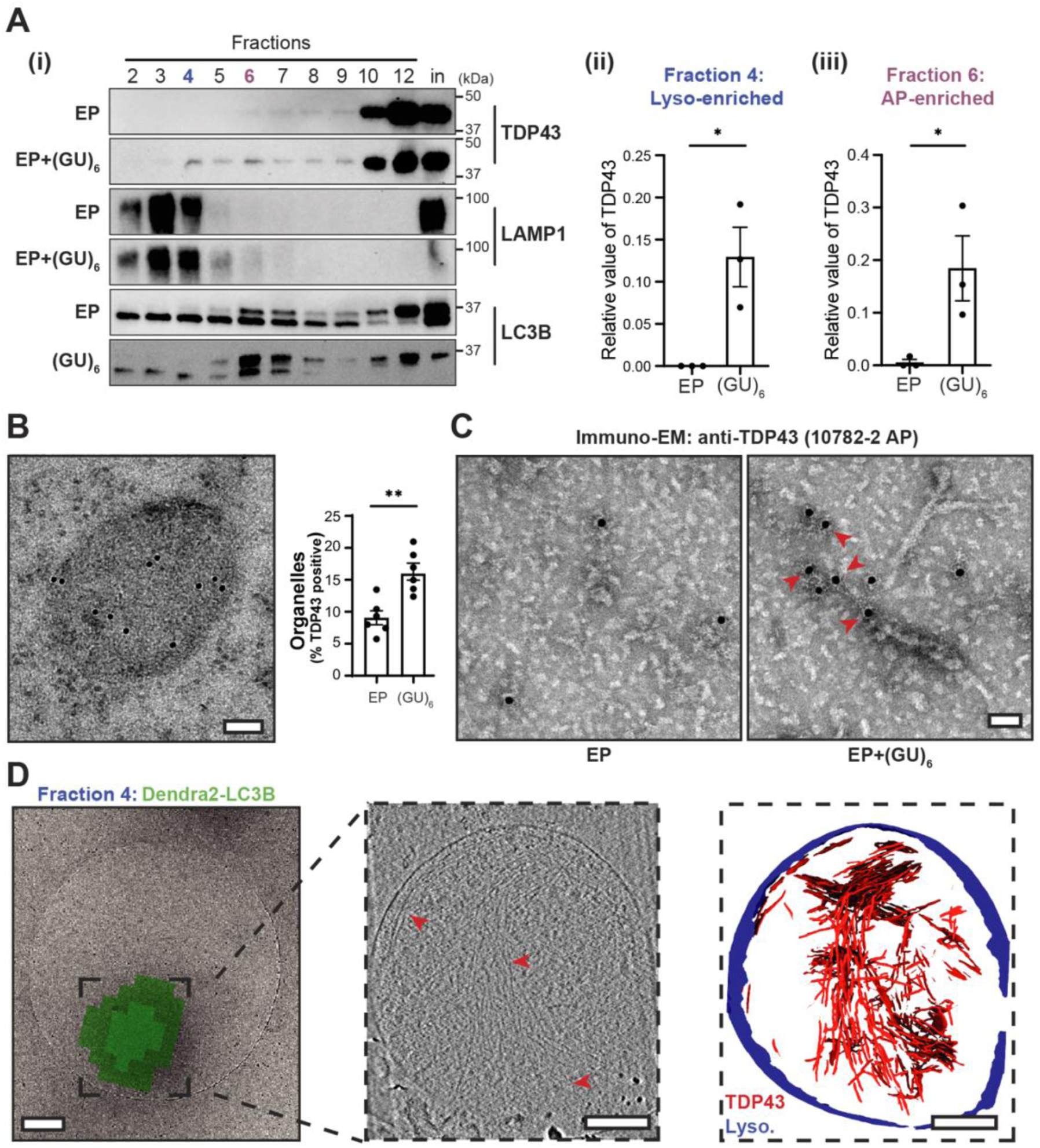
GU-rich oligonucleotides induce TDP43 fibrilization in autophagosomes and lysosomes of HEK293T cells. (**A**) (i) Representative immunoblots of fractions 2-12 and membrane fraction input (in) isolated from Endo-Porter + (GU)_6_-treated Dendra2-LC3B HEK293T cells, detected with anti-TDP43, -LAMP1, and -LC3B antibodies. n=3. Quantification of TDP43 in fractions 4 (ii) and 6 (iii) (mean 土 SEM, Tukey test, *p < 0.05). (**B**) Anti-TDP43 immunogold labeling on an ultrathin section of (GU)_6_-treated Dendra2-LC3B HEK293T cells (left). n=6. Scale bar, 100nm. Quantification of TDP43-positive vesicles in oligo-treated Dendra2-LC3B HEK293T cells (right). (Welch’s t-test, **p < 0.01). (**C**) Representative negative stain EM images of anti-TDP43 immunogold labeled, detergent-insoluble TDP43 fibrils isolated from Dendra2-LC3B HEK293T cells. Red arrowheads depict gold beads on fibrils. Scale bar, 50nm. (**D**) Correlative cryo-electron tomography of lysosome-enriched fraction 4 from (GU)_6_-treated Dendra2-LC3B HEK293T cells. A representative 6500× cryo-TEM montage image of a lysosome overlaid with the Dendra2-LC3B cryo-fluorescence signal (left). Slice through a reconstructed, denoised tomogram of the same region, highlighting ensconced TDP43 fibrils (red arrowheads, middle). Segmentation of the tomogram with TDP43 fibrils (red) and the lysosome membrane (blue) highlighted (right). Scale bars: 200nm.

To confirm this, we prepared ultrathin sections of high-pressure frozen and freeze-substituted Dendra2-LC3B HEK293T cells, allowing us to examine RNA-induced TDP43 mislocalization by immuno-electron microscopy (immuno-EM). Gold-labeled anti-TDP43 antibodies reaffirmed the presence of TDP43 within single membrane-bound lysosomes (**Figure 4B).** Membrane-bound vesicles exhibited significantly more gold beads in (GU)_6_-treated cells in comparison to cells treated with Endo-Porter alone (control), indicative of greater lysosomal TDP43 immunoreactivity following RNA application (**Figures 4B and S7**).

We also isolated sarkosyl-insoluble fractions from control and (GU)_6_-treated Dendra2-LC3B HEK293T cells to assess for TDP43 fibrils analogous to those detected in iNeurons (**Figure S6B**). Negative stain transmission electron microscopy (TEM) of these samples revealed the presence of fibrils, reminiscent of TDP43 filaments obtained from post-mortem brain tissue^28,29^, but only in samples from (GU)_6_-treated cells. Immunogold labeling with TDP43-specific antibodies once again confirmed that these fibrils are comprised of TDP43 (**Figure 4C**). To determine if fibrils are already present in isolated organelles, and to verify that they are not an artifact of biochemical extraction, we performed cryo-CLEM on plunge-frozen autophagosomal and lysosomal fractions isolated from Dendra2-LC3B HEK293T cells treated with (GU)_6_ oligos (**Figures 4D and S6C**). In these samples, Dendra2-LC3B fluorescence enabled the precise localization of autophagosomes and lysosomes. Subsequent cryo-ET tilt series data collection from these areas, followed by tomogram reconstruction, revealed multiple filaments ensconced within membrane-bound organelles (red arrows, **Figures 4D and S6C**). These filaments displayed a mean width of 4.9 ± 0.79nm, mirroring both iNeuron intralysosomal filaments and the amyloidogenic C-terminus of TDP43 obtained by limited proteolysis of the protein^61^ (**Figure 3E**).

### TDP43 mislocalizes to autophagosomes and lysosomes in ALS patient tissue

Our observations of TDP43 fibrils within autophagosomes and lysosomes suggest that the earliest stages of pathological TDP43 deposition may occur within these organelles. To pursue this possibility, we performed dual immunohistochemistry for TDP43 and markers of lysosomes (LAMP1) or autophagosomes (LC3B) in ALS patient spinal cord sections. Within the patient tissue, we stratified ventral motor neurons based on TDP43 localization. Motor neurons with strong nuclear TDP43 signal exhibited the expected distribution of LAMP1 and LC3B within the cytosol (**Figure 5A**), as did motor neurons from non-ALS controls (**Figures S8A**). However, ALS motor neurons with nuclear TDP43 exclusion displayed a marked overlap of cytoplasmic TDP43 puncta with lysosomes, and to a lesser extent, autophagosomes (**Figure 5A**). The overlap between TDP43 and these organelles was further confirmed by immunofluorescence microscopy in neurons displaying TDP43 mislocalization, but not in ALS neurons with nuclear TDP43 or neurons from non-ALS controls (**Figures 5B, S8B**). Quantification of control and ALS patient tissue confirmed a significant increase in TDP43 colocalization with LAMP1 in neurons with nuclear-excluded TDP43 (**Figure 5C**). On the contrary, colocalization of TDP43 with the autophagosomal marker LC3B was greater in control neurons than in ALS neurons (**Figure 5D**), suggesting that TDP43 may be actively cleared or eliminated by these organelles.

**Figure 5.**
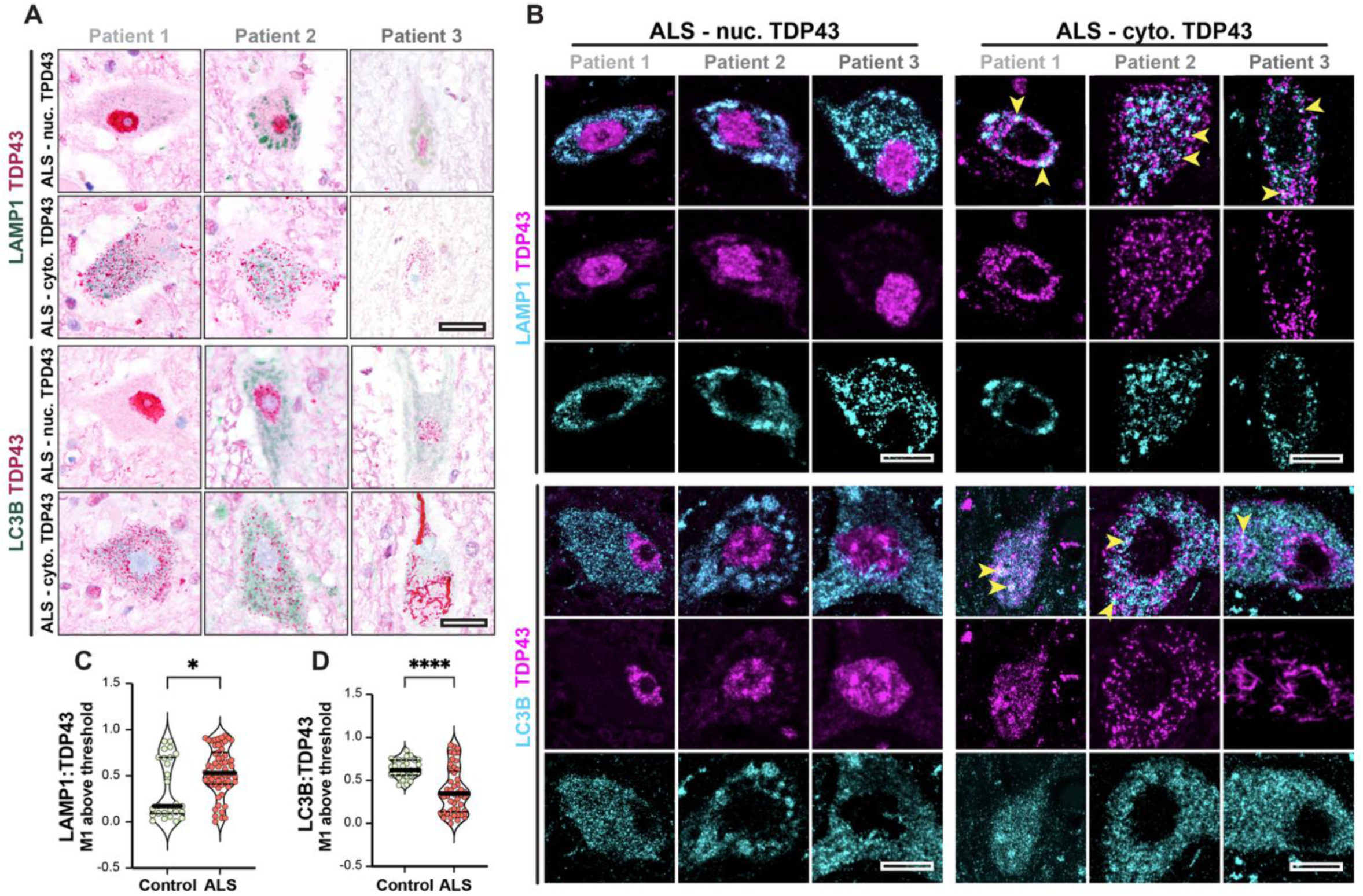
TDP43 deposits overlap with lysosomal markers in patient tissue. (**A**) Immunohistochemical analysis of motor neurons from the spinal cords of three patients (1, 2, 3) with sporadic ALS, showing either TDP43 (magenta) and LAMP1 (green, top) or LC3B (green, bottom). In each case, motor neurons with nuclear TDP43 (‘ALS–nuc. TDP43’, top row) and neurons with cytoplasmic TDP43 (‘ALS-cyto. TDP43’, bottom row) are shown. Scale bar, 20µm. (**B**) Representative immunofluorescence images of ALS spinal motor neurons (from patients 1, 2, and 3) stained for TDP43 (magenta), LAMP1 (cyan, top half), or LC3B (cyan, bottom half). Motor neurons with nuclear TDP43 are to the left, while those with cytoplasmic TDP43 are to the right. Colocalization of LAMP1 or LC3B with cytosolic TDP43 marked by yellow arrowheads. Scale bars, 10 µm. **(C)** Proportion of LAMP1 fluorescence overlapping with TDP43 in non-ALS control and ALS motor neurons. *p < 0.05, Mann-Whitney test; n=26 and n=60 motor neurons across 3 control and 3 ALS patients, respectively. **(D)** Proportion of LC3B fluorescence overlapping with TDP43 in 3 separate control and ALS tissue sections. ****p < 0.0001, Mann-Whitney test; n=29 and n=48 motor neurons across 3 control and 3 ALS patients, respectively. In (C) and (D), median and quartiles are shown as thick and dashed lines, respectively.

To verify the accumulation of mislocalized TPD43 within autophagosomes and lysosomes of post-mortem patient material, we adapted the method we previously used for isolating these organelles from cultured cells and used it to fractionate ALS/FTLD-TDP prefrontal cortex (**Figures 6A and S6A**). In lysosome-rich fractions, we detected full-length and pTDP43 (S409/S410) as well as truncated species of ∼35 and 25 kDa (**Figure 6B**). Cryo-ET analysis of these lysosomes revealed filamentous inclusions within membrane-bound structures that were remarkably similar to what we observed in iNeurons and HEK293T cells (**Figure 6C**). The widths of filaments in patient-derived lysosomes displayed a bimodal distribution, ranging from 5 to 15nm (**Figures 6D, E**). These filaments most likely represent distinct, partially proteolyzed states of TDP43. Finally, we extracted sarkosyl-insoluble material from patient-derived brain lysosomes (**Figure S6B**) and probed for pTDP43 (S409/S410) by immuno-EM. We observed filaments decorated with gold beads, confirming the presence of TDP43-positive fibrillar deposits within the lysosomes of human patient brain tissue (**Figure 6F**). Together, these findings validate and extend our observations from (GU)_6_-treated iNeurons, suggesting that RNA dysregulation may similarly foster TDP43 mislocalization and aggregation in disease.

**Figure 6.**
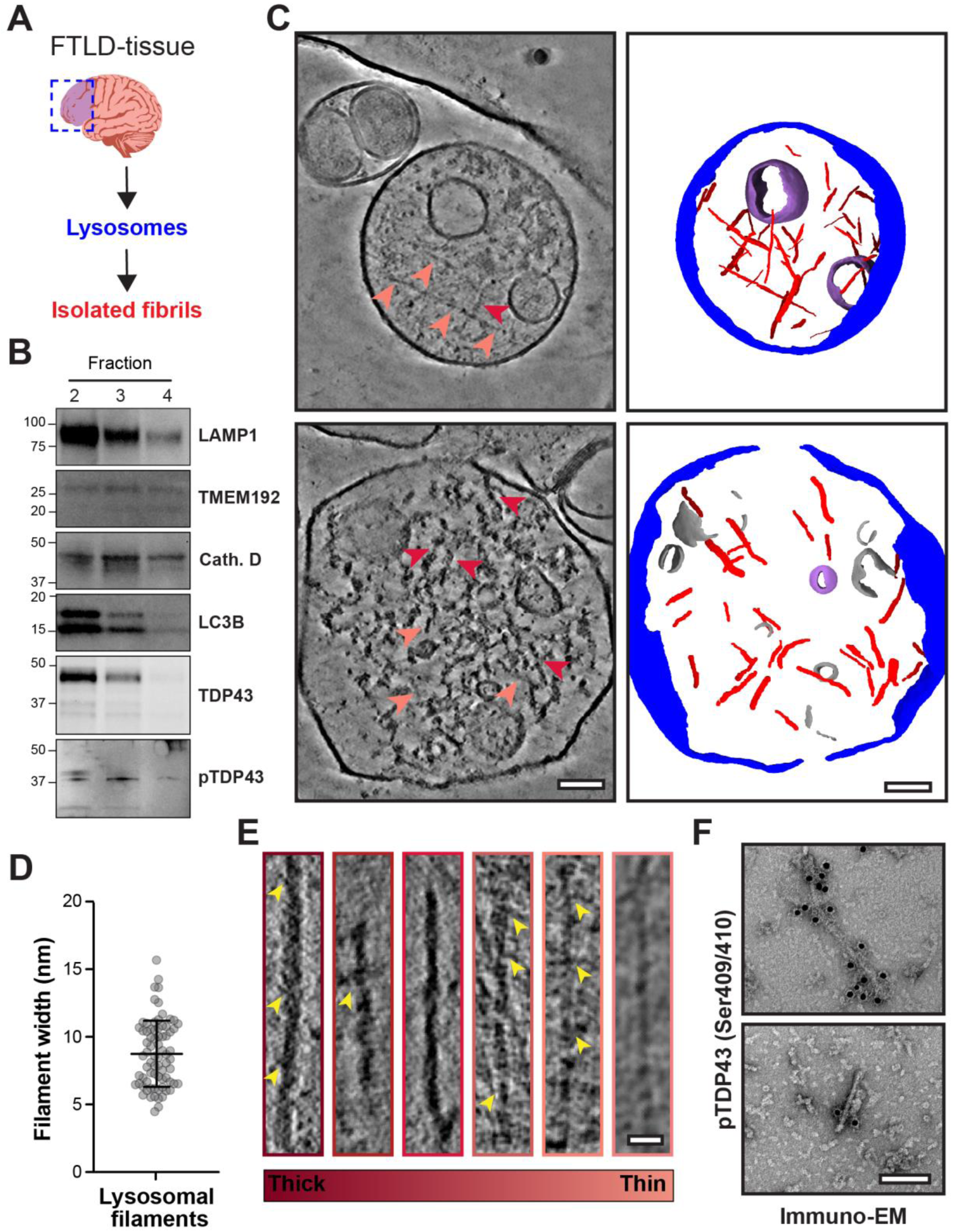
Patient brain-derived lysosomal TDP43. **(A)** Schematic of post-mortem brain tissue serial ultracentrifugation and density gradient fractionation (see Figure S6 for more details). (**B**) Immunoblots of fractions 2-4 from the density gradient fractionation of ALS/FTLD-TDP brain tissue probed for the lysosomal markers LAMP1, TMEM192, and Cathepsin D (Cath. D); autophagosomal marker, LC3B; TDP43 and phospho-TDP43 (pTDP43, S409/410). TDP43 is enriched in autophagosomes and lysosomes, specifically in fraction 2 and 3, confirming the presence of multiple proteolyzed TDP43 species. (**C**) Left: slice through two reconstructed and denoised cryo-tomograms of fraction F3, revealing filaments of varying widths and morphology within single-membraned lysosomes. Arrowheads depict filaments of varying widths. Thicker ones are represented in red, while the thinner ones are in light red. Scale bar: 50nm. Right: Corresponding segmentation and rendering of the tomogram on the left showing the lysosomal membrane (blue), TDP43 fibrils (red), unknown membranous cargo (gray), and vesicles (lilac). (**D**) Plot of lysosome-bound filament widths ranging from ∼5 to ∼15nm, analyzed from 21 tomograms. (**E**) Gallery of TDP43 filaments within the lysosomes. Slice through the tomograms cropped to show filaments of varying thickness. Yellow arrowheads indicate additional densities branching out of the filaments. Scale bar: 20nm. (**F**) Immunogold labeling against phosphorylated TDP43 (pTDP43, Ser409/410) on sarkosyl-insoluble filaments isolated from the lysosome-enriched fraction (F3). Scale bar: 50nm.

## Discussion

TDP43 recognizes nearly one-third of all transcribed genes and is crucial for diverse aspects of RNA metabolism^62,63^. Loss of this protein from the nucleus and its cytosolic accumulation are closely tied to neurodegeneration in cellular and animal models of ALS/FTLD-TDP, as well as in human post-mortem samples^14,54,64^. Indeed, TDP43 knockdown leads to dramatic and widespread changes in RNA splicing and stability^25,44,45,65–67^, implying that RNA misprocessing in ALS/FTLD-TDP is downstream of TDP43 mislocalization. Our data, together with recent studies highlighting the significance of RNA for TDP43 localization^22–24^, are consistent with an alternative possibility: RNA dyshomeostasis may precede and drive TDP43 mislocalization, thereby eliciting a self-perpetuating positive feedback loop that further facilitates and enhances RNA misprocessing. The result is a spiral of RNA dysfunction and TDP43 mislocalization that expedites neuronal toxicity and degeneration in disease.

RNA misprocessing may originate from genetic and/or acquired means. For instance, mutations in genes encoding RNA-binding proteins, such as *FUS*, *HNRNPA2B1*, *MATR3*, and *TARDBP*, themselves lead to ALS and FTLD-TDP^68–74^. Co-transcriptional RNA modifications, including methylation at the N^6^ position of adenosine (m^6^A), may also affect TDP43 binding affinity and/or the fate of transcribed RNA^26,75–78^. Abnormal RNA methylation is characteristic not only of ALS and FTLD-TDP but also of other conditions, such as Alzheimer’s disease, that display TDP43 pathology^26,27,79–81^. RNA-based sequestration of TDP43 into cytosolic granules is also observed during skeletal muscle regeneration in mice and humans^82^. These myo-granules are rich in fibrillar TDP43, suggesting common misfolding events upon cytosolic TDP43 mislocalization. Here, we utilized an idealized TDP43 substrate—GU-rich RNA oligos—to initiate the efflux of TDP43 from the nucleus in iNeurons (**Figures 1A-D and S1A-F**). Treated iNeurons accumulate cryptic isoforms from TDP43-targeted transcripts and display progressive toxicity over time (**Figures 1E and 1G**), effectively recapitulating key hallmarks of the pathophysiology of ALS/FTLD.

Biochemical fractionation experiments revealed the presence of full-length TDP43 within autophagosomes and lysosomes of (GU)_6_-treated cells (**Figure 4**). This is consistent with the appearance of full-length TDP43 within pathological aggregates in ALS spinal motor neurons^12^, the overlap between TDP43 inclusions and lysosomal markers in (GU)_6_-treated iNeurons and post-mortem ALS samples (**Figures S3A and S3D; Figures 5B-D**), the visualization of TDP43 filaments within autophagosomes and lysosomes in iNeurons (**Figures 3A-C**), and the appearance of filaments in purified lysosomes from ALS/FTLD-TDP brain (**Figure 6C**). Collectively, these data imply that mislocalized TDP43 accumulates and misfolds within acidified organelles. Future investigations may focus on whether the fibrils represent pathological intermediates capable of seeding further TDP43 aggregation, as well as the factors governing the formation and/or degradation of such intermediates.

Structural studies on purified protein fragments or fibrils extracted from patient samples are invaluable for deriving small-molecule ligands or tracers specific for protein inclusions; nevertheless, these investigations are limited in their ability to uncover molecular mechanisms of TDP43 aggregation. To address this, we established an *in situ* cryo-CLEM platform (**Figure 2**), which provides the first visual snapshots of endogenous TDP43 mislocalization and aggregation within human neurons. Application of short GU-rich oligos leads to the appearance of ‘exclusion-zones’ containing TDP43 inclusions within the cytosol (**Figure 3A, Video S2**). The material properties of the inclusions or condensates (**Figure 3A**, solid-like vs. amorphous) may depend on chaperones (i.e., HSPB1)^83^ and/or RNA length, size, modifications, and motif valency^84^. Overexpression of the intrinsically aggregation-prone, GFP-tagged C-terminal fragment of TDP-43 (GFP–TDP-25) leads to the formation of ‘gel-like’, amorphous cytoplasmic inclusions^85^. Such experimental conditions, while helpful in probing structures formed by TDP43 fragments, may artificially accelerate or bias the underlying mechanisms of assembly and thus limit the physiological relevance of the conclusions. In contrast, our study demonstrates TDP43 fibrilization at endogenous protein levels, thereby providing valuable insights under native expression conditions that more faithfully reflect the physiological and pathological context.

Cytosolic TDP43 inclusions were often found in association with lysosomal membranes, but within these organelles, TDP43 adopts filamentous structures (**Figure 3A and C; and Videos S2 and S4**). The acidified environment of lysosomes may promote the further demixing of TDP43 from condensates, thereby enhancing its misfolding and promoting the formation of large, ordered filaments^86–88^. The appearance of TDP43 within autophagosomes (**Figure 3B, and Video S3**) argues against chaperone-mediated autophagy in the clearance of cytosolic TDP43^89^ but would be consistent with bulk macroautophagy. Additionally, we cannot rule out the possibility of selective autophagy of fibrillar TDP43^90,91^, involving an as-yet-unidentified adapter. Prior studies identified a single LC3-interacting region (LIR)-containing protein in association with TDP43— TATA-box binding protein (TBP)^92^—but further studies are required to determine whether this factor is necessary for the delivery of cytosolic TDP43 to lysosomes and/or autophagosomes.

The TDP43 C-terminal low-complexity domain (LCD) is critical for forming pathological amyloid-like aggregates *in vitro*^9,10^. Within this region, amino acid stretches 286-331 in primary cortical neurons and 341-367 in neuronal cell lines are sufficient for fibril formation^93^. Here, we show that introducing GU-rich oligos reproduces TDP43 mislocalization and aggregation within human neurons and cell lines. Mislocalized TDP43 forms structures reminiscent of patient-derived fibrils in autophagosomes and lysosomes, underscoring the importance of acidified organelles in the formation and/or clearance of nascent TDP43 aggregates (**Figure 3**). The architecture of TDP43 species within lysosomal/autophagosomal fibrils, and whether these structures sequester specific RNAs or proteins, remains to be investigated.

Based on these observations, we propose a model for TDP43 mislocalization, clearance, and its aggregation in disease (**Figure 7**). External stressors (i.e., injury) or internal events (i.e., genetic mutations, RNA dyshomeostasis) may trigger nuclear TDP43 egress (**Figure 7A**). In the cytosol, TDP43 undergoes liquid-like condensation, as observed for other RNA-binding proteins in similar environments^94^. At this stage, at least two scenarios are possible. If the stressors are self-limiting and autophagy is functional, mislocalized TDP43 is eliminated, and nuclear TDP43 levels are restored through new protein synthesis. Conversely, if the stressors persist, both existing and newly synthesized pools of TDP43 are mislocalized, accelerating TDP43 accumulation within autophagosomes and lysosomes (**Figures 7B and 7C**)^95^.

**Figure 7.**
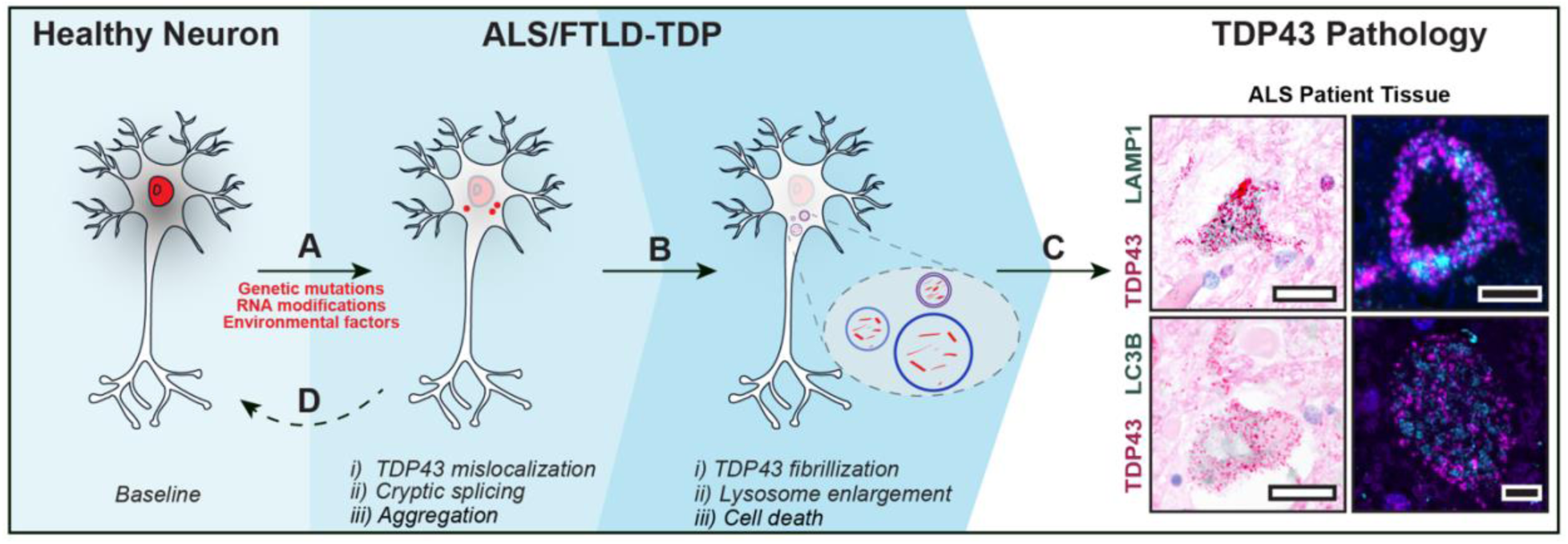
An autophagy-centric model of early-stage TDP43 pathology. (**A**) TDP43 (red) primarily localizes to the nucleus in healthy neurons. RNA misprocessing, genetic mutations, or environmental factors may trigger the nuclear egress and cytosolic accumulation of TDP43. TDP43 nuclear egress leads to cryptic splicing and the formation of cytosolic TDP43 aggregates. (**B**) Mislocalized TDP43 accumulates in autophagosomes and lysosomes, resulting in their enlargement. (**C**) Continued TDP43 mislocalization enhances fibrilization and neuronal loss, as observed in ALS, FTLD-TDP, and other TDP43 proteinopathies. Representative immunohistochemistry (left) and immunofluorescence (right) images of ALS motor neurons stained for TDP43, LAMP1 (lysosomes, top row), and LC3B (autophagosomes, bottom row). Scale bars: 20µm (left) and 5µm (right). (**D**) Restoring TDP43 nuclear localization at initial stages may reverse molecular hallmarks of disease and forestall neurodegeneration.

Our data reveal a size-dependent relationship between lysosomes and intraluminal fibrils, with larger fibrils preferentially residing in enlarged lysosomes — suggesting impaired lysosomal processing or lysosomal dysfunction (**Figures 3D and 3E**)^95,96^. The precise characteristics and turnover capacity of enlarged lysosomes containing large TDP43 fibrils remain to be determined; however, their identification with LysoTracker strongly suggests that they retain an acidic pH. Alternatively, fibrils may nucleate in cytosolic condensates, followed by internalization into lysosomes via an unknown receptor. Subsequent lysosomal fusion and volumetric expansion may facilitate continued fibril elongation, leading to the accumulation of larger filamentous aggregates. As is the case for α-synuclein^97^, TDP43 fibrils may escape from lysosomes and seed additional TDP43 aggregation within the cytoplasm. Lysosomal TDP43 may also be delivered to autophagosomes, where it undergoes either degradative or secretory autophagy; acquired or genetic perturbations may further enhance the latter event, thereby increasing lysosomal/autophagosomal activity^98^. These possibilities are consistent with disease-associated mutations in autophagy-related genes (i.e., *UBQLN2, C9ORF72, OPTN, SQSTM1*)^99–103^, genes involved in lysosomal trafficking and function (*GRN, CHMP2B*)^104–109^, age-related declines in the efficiency of neuronal autophagy, and the broad neuroprotection afforded by autophagy-inducing strategies in ALS/FTLD-TDP models^40,53,54^.

Taken together, we establish a cellular model of TDP43 mislocalization and cytoplasmic aggregation, a signature pathologic event in ALS/FTLD-TDP. Using this model, we provide the first *in situ* visual snapshots of TDP43 aggregates within human neuronal lysosomes and autophagosomes, highlighting the critical connection between autophagy and the pathogenesis of ALS/FTLD-TDP. We expect this platform to be invaluable not only for investigating the pathogenesis and origins of TDP43 pathology but also for developing strategies to detect and potentially even reverse TDP43 mislocalization in ALS/FTLD-TDP and related TDP43-proteinopathies.

## Acknowledgments

We thank all members of the Barmada, Baldridge, and Mosalaganti laboratories for their suggestions. We thank Dr. Jing Liang and the Microscopy Core of the University of Michigan Biomedical Research Core Facilities for their assistance with high-pressure freezing, freeze-substitution, and resin embedding of HEK293T cells. We thank Dr. Vinson Lam for his assistance with cryo-ET data collection. We thank Dr. Luke Lavis at the Janelia Research Campus for sharing JF635. This work was supported by: National Institutes of Health DP2GM150019-01, and Klatskin Sutker Discovery Fund (S.M.), National Institutes of Health R01NS097542, R01NS113943 and 1R56NS128110-01 (S.J.B), Kissick Family Foundation & Milken Institute Frontotemporal Dementia Grant (S.J.B. & S.M.), University of Michigan Pioneer Postdoctoral Fellowship (A.L.E.), University of Michigan Life Sciences Institute, American Heart Association Predoctoral Fellowship (R.S.), National Institutes of Health AWD012778 (E.P.), National Institutes of Health R35GM128592 (R.D.B.), National Institutes of Health T32GM007544 (J.E.R.), National Institutes of Health T32GM141840 to (M.G.F.), National Institutes of Health P30AG072931 to the University of Michigan Brain Bank and Alzheimer’s Disease Research Center, National Institutes of Health S10OD030275 and the Arnold and Mabel Beckman Foundation award to the University of Michigan Cryo-EM facility. The funders had no role in the study design, data collection, analysis, or the content and publication of this manuscript.

## Author contributions

A.L.E. prepared, collected, and analyzed the WT and HaloTag-TDP43 iNeuron cryo-CLEM/cryo-ET data. M.G.F. prepared patient tissue derived lysosomes and sarkosyl-insoluble filaments, and collected and analyzed immunoblots, immuno-EM, and cryo-ET data. A.L.E., M.L.C., and J.E.R. prepared HEK293T lysosome samples; M.L.C. performed and analyzed immunoblots and immunogold labeling experiments, while M.G.F. collected and analyzed cryo-CLEM and cryo-ET data; A.L.E., M.L.C., and R.S. performed segmentation analysis. D.A. prepared the iNeuron samples and collected the iNeuron confocal microscopy and immunoblotting data; A.L.E., M.G.F., and X.S. analyzed the iNeuron confocal microscopy data. D.A. prepared the iNeuron samples for qPCR analysis; M.B. collected and analyzed qPCR data and survival assays. E.S.P. and D.T. prepared and collected patient tissue duplex immunohistochemistry and immunofluorescence data; A.E. and X.S. analyzed patient tissue immunohistochemistry and immunofluorescence data. M.G.F. wrote custom scripts for aligning and reconstructing tilt series. E.M.H.T. generated and maintained all the iPSC lines. A.L.E., S.J.B., and S.M. conceptualized the study, supervised the project, and wrote the manuscript.

## Resource availability

### Lead contact

Please direct the requests for resources and reagents to the lead contact, Shyamal Mosalaganti.

### Materials availability

Materials are available upon reasonable request to the lead contact. Additional details are provided in the materials and methods section.

## Figures (Main, 1-7)

Figure 1. Short GU-rich oligonucleotides induce TDP43 nuclear export, cryptic exon splicing, and neurotoxicity in human iNeurons.

Figure 2. *In situ* cryo-CLEM workflow for ultrastructural analysis of human iNeurons.

Figure 3. Mislocalized TDP43 accumulates within neuronal autophagosomes and lysosomes.

Figure 4. GU-rich oligonucleotides induce TDP43 fibrilization in autophagosomes and lysosomes of HEK293T cells.

Figure 5. TDP43 deposits overlap with lysosomal markers in patient tissue.

Figure 6. Patient brain-derived lysosomal TDP43.

Figure 7. An autophagy-centric model of early-stage TDP43 pathology.

## Figures (Supporting, S1-S8)

Figure S1. TDP43 nuclear egress and neurotoxicity in human iNeurons treated with short GU-rich oligonucleotides.

Figure S2. Prolonged treatment of iNeurons with GU-rich oligonucleotides induces TDP43 phosphorylation (S409/410).

Figure S3. TDP43 colocalizes with lysosomes in human iNeurons treated with short GU-rich oligonucleotides.

Figure S4. HaloTag-TDP43 iNeurons for *in situ* cryo-ET analysis.

Figure S5. Gallery of lysosomes and autophagosomes from Halo-TDP43 iNeurons.

Figure S6. Isolation of autophagosomes and lysosomes from (GU)_6_-treated HEK293T cells or ALS/FTLD patient postmortem brain tissue.

Figure S7. Immunogold labelling of HEK293T cell sections.

Figure S8. TDP43 and lysosome/autophagosome staining in control patient tissue.

**Figure. S1:**
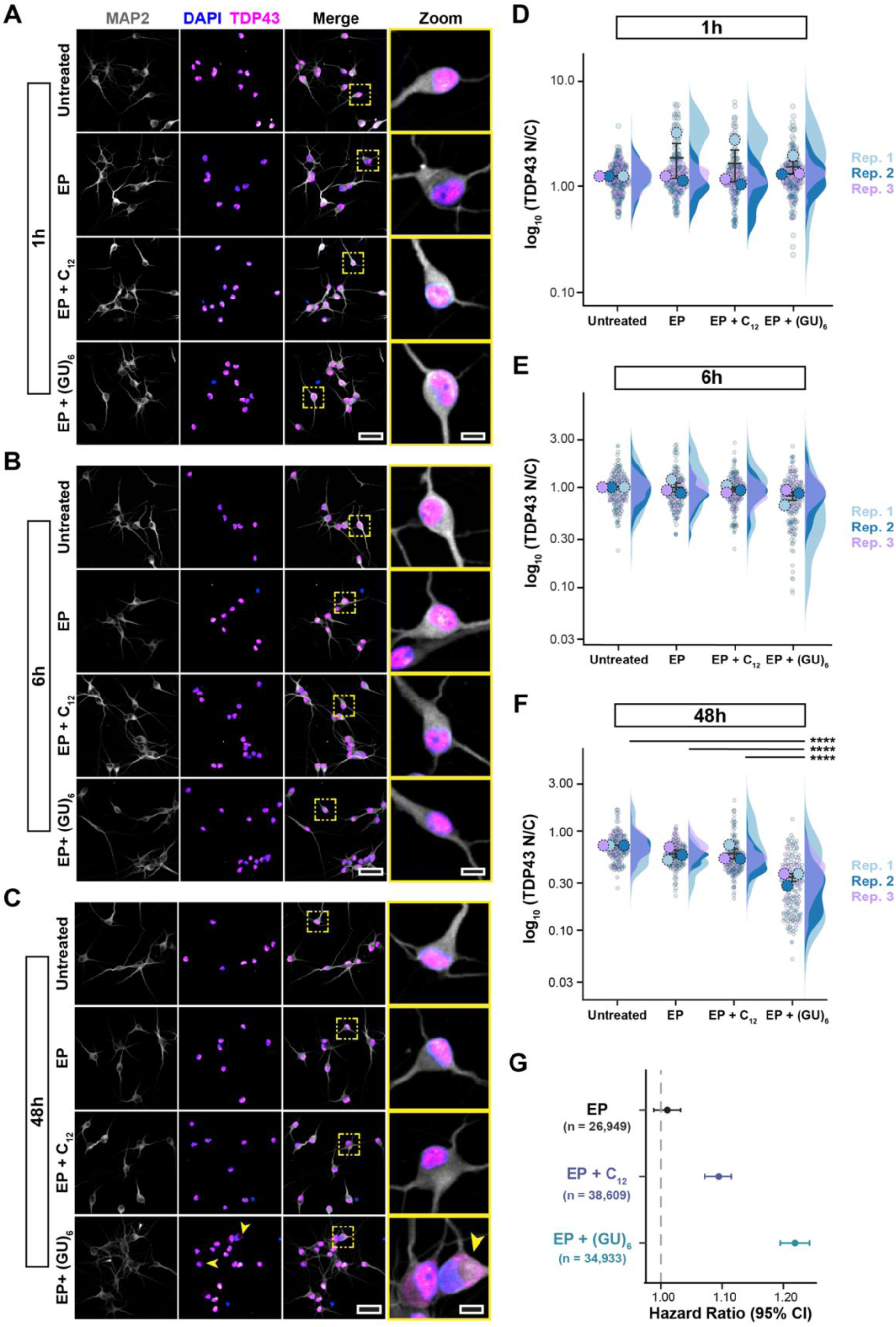
TDP43 nuclear egress and neurotoxicity in human iNeurons treated with short GU-rich oligonucleotides. (**A-C**) Representative confocal fluorescence microscopy images of untreated DIV10 iNeurons (top row) or iNeurons treated with EP only (second row), EP with 2µM C_12_ (third row), and EP and 2 µM (GU)_6_ (last row) for 1h (A), 6h (B), and 48h (C) before being chemically fixed and stained for neuronal marker (MAP2), nuclei marker (DAPI), and TDP43. TDP43 mislocalization from the nucleus to cytoplasm was evident in (GU)_6_-treated intact iNeurons treated for longer than 24h (C, last row); yellow arrowheads highlight select iNeurons. Representative iNeurons (yellow dashed box) for each condition are shown at 5× magnification (last column). Images are pseudo-colored as follows: MAP2 (gray), DAPI (blue), and TDP43 (magenta). Scale bars: 50µm, except for the 5× magnified images, which are 10µm. (**D-F**) Log-transformed ratio of TDP43 fluorescence intensity in the nucleus vs. cytoplasm (N/C ratio) plotted for DIV10 iNeurons either untreated or treated with EP only, EP + 2µM C_12_, or EP + 2µM (GU)_6_ for 1h (D), 6h (E), or 48h (F). TDP43 N/C ratios were normalized to the untreated condition for each replicate. (mean 土 SEM, Kruskal-Wallis test followed by Dunn’s post hoc test, ****p < 0.0001, ***p<0.001, **p<0.01, *p<0.05; n > 140 across 3 biological replicates). Biological replicates 1-3 are colored as light blue, blue, and purple, respectively. (**G**) Forest plot of hazard ratios ± 95%CI for human iNeurons treated with EP alone (black), EP + C_12_ (purple), or EP + (GU)_6_ oligos (teal) (2µM each). PBS serves as a reference (hazard ratio = 1). n = number of iNeurons, combined from 6 technical replicates for each of the 3 biological replicates. * Hazard ratio (HR) = 1.09, p <2×10^-16^; ** HR= 1.22, p <2×10^-16^; Cox proportional hazards analysis.

**Figure S2.**
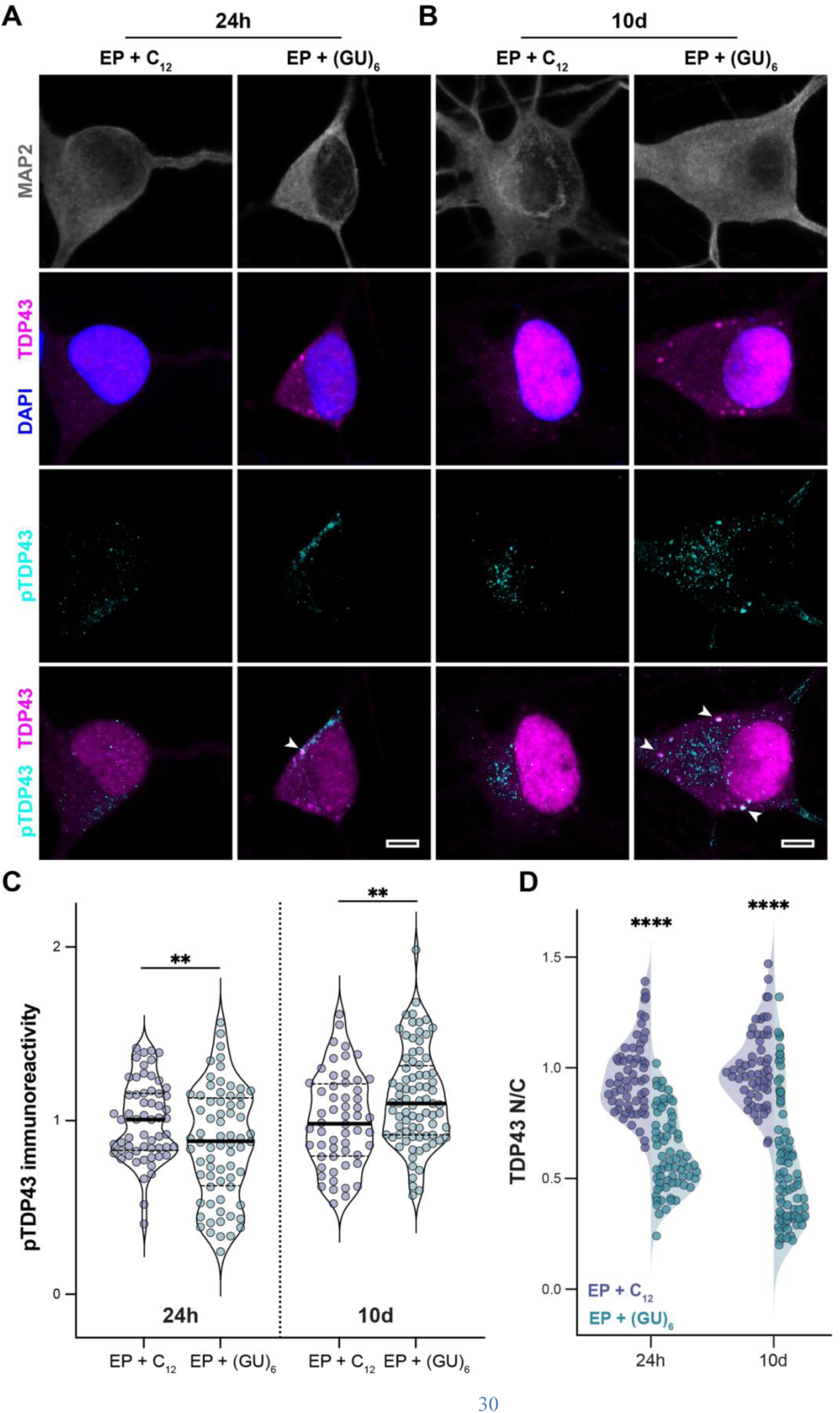
Prolonged treatment of iNeurons with GU-rich oligonucleotides induces TDP43 phosphorylation (S409/410). (**A-B**) Representative confocal fluorescence microscopy images of DIV10 iNeurons treated with EP + C_12_ or EP + (GU)_6_ (2µM each) for 24h (A) and 10d (B) prior to immunostaining for MAP2 (grey), TDP43 (magenta), and pTDP43 (S409/410, cyan). Scale bar: 5µm. White arrowheads highlight TDP43 punctae colocalizing with pTDP43. (**C**) pTDP43 mean fluorescence intensity plotted for DIV10 iNeurons treated with EP + C_12_ or EP + (GU)_6_ (2µM each) for 24h or 10d. pTDP43 immunoreactivity values were normalized to the EP + C_12_ condition for each replicate (Welch’s t-test, **p <0.01, n > 56 across three biological replicates). Median and quartiles are shown as thick and dashed lines, respectively. (**D**) Ratio of TDP43 fluorescence intensity in the nucleus vs. cytoplasm (N/C ratio) plotted for DIV10 iNeurons treated with EP + C_12_ or EP + (GU)_6_ (2µM each) for 24h or 10d. TDP43 N/C ratios were normalized to the EP + C_12_ condition for each replicate (Mann-Whitney test, ****p < 0.0001, n > 56 across three biological replicates).

**Figure S3.**
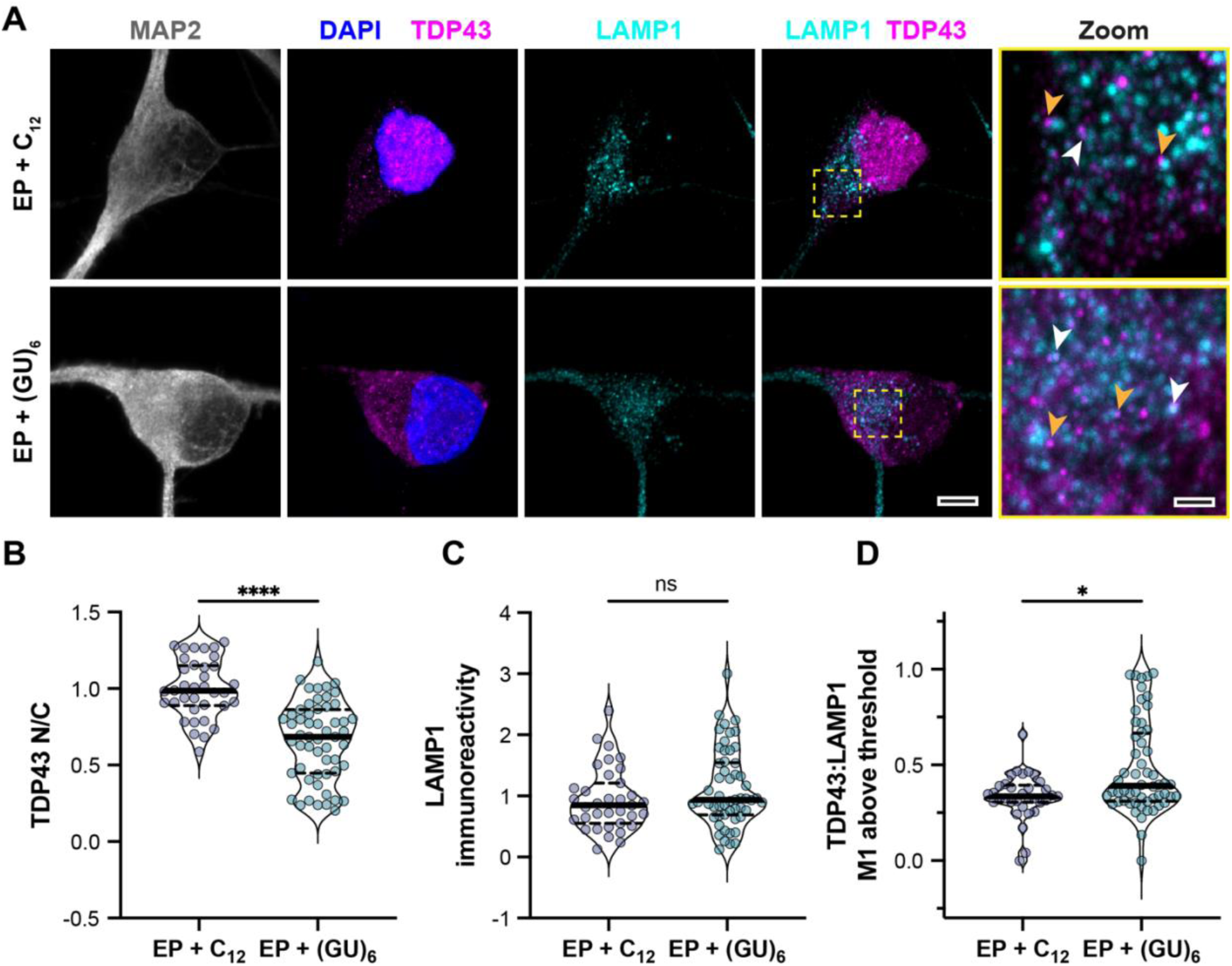
TDP43 colocalizes with lysosomes in human iNeurons treated with short GU-rich oligonucleotides. (**A**) Representative confocal fluorescence microscopy images of DIV10 iNeurons treated with EP + C_12_ or EP + (GU)_6_ (2µM each) for 24h prior to immunostaining for MAP2 (grey), TDP43 (magenta), and LAMP1 (cyan). The region of the iNeuron soma highlighted by the dashed yellow box is 5× magnified (last column). White arrowheads highlight TDP43 punctae colocalizing with lysosomes. Orange arrowheads highlight TDP43 punctae adjacent to lysosomes. Scale bar: 5µm, except for the 5× magnified image, which is 1µm. (**B**) Ratio of TDP43 fluorescence intensity in the nucleus vs. cytoplasm (N/C ratio) plotted for DIV10 iNeurons treated with EP + C_12_ or EP + (GU)_6_ (2µM each) for 24 h. TDP43 N/C ratios values were normalized to the EP + 2µM C_12_ condition for each replicate (Mann-Whitney test, ****p < 0.0001, n > 36 across three biological replicates). (**C**) LAMP1 mean fluorescence intensity plotted for DIV10 iNeurons treated with EP + C_12_ or EP + (GU)_6_ (2µM each) for 24 h. LAMP1 immunoreactivity values were normalized to the EP + 2µM C_12_ condition for each replicate (Mann-Whitney test, n > 36 across three biological replicates). (**D**) Proportion of LAMP1 fluorescence overlapping with TDP43 in DIV10 iNeurons treated EP + C_12_or EP + (GU)6 (2µM each) for 24h (Mann-Whitney test, *p < 0.05, n>33 across three replicates). Median and quartiles for (B-D) are shown as thick and dashed lines, respectively.

**Figure S4.**
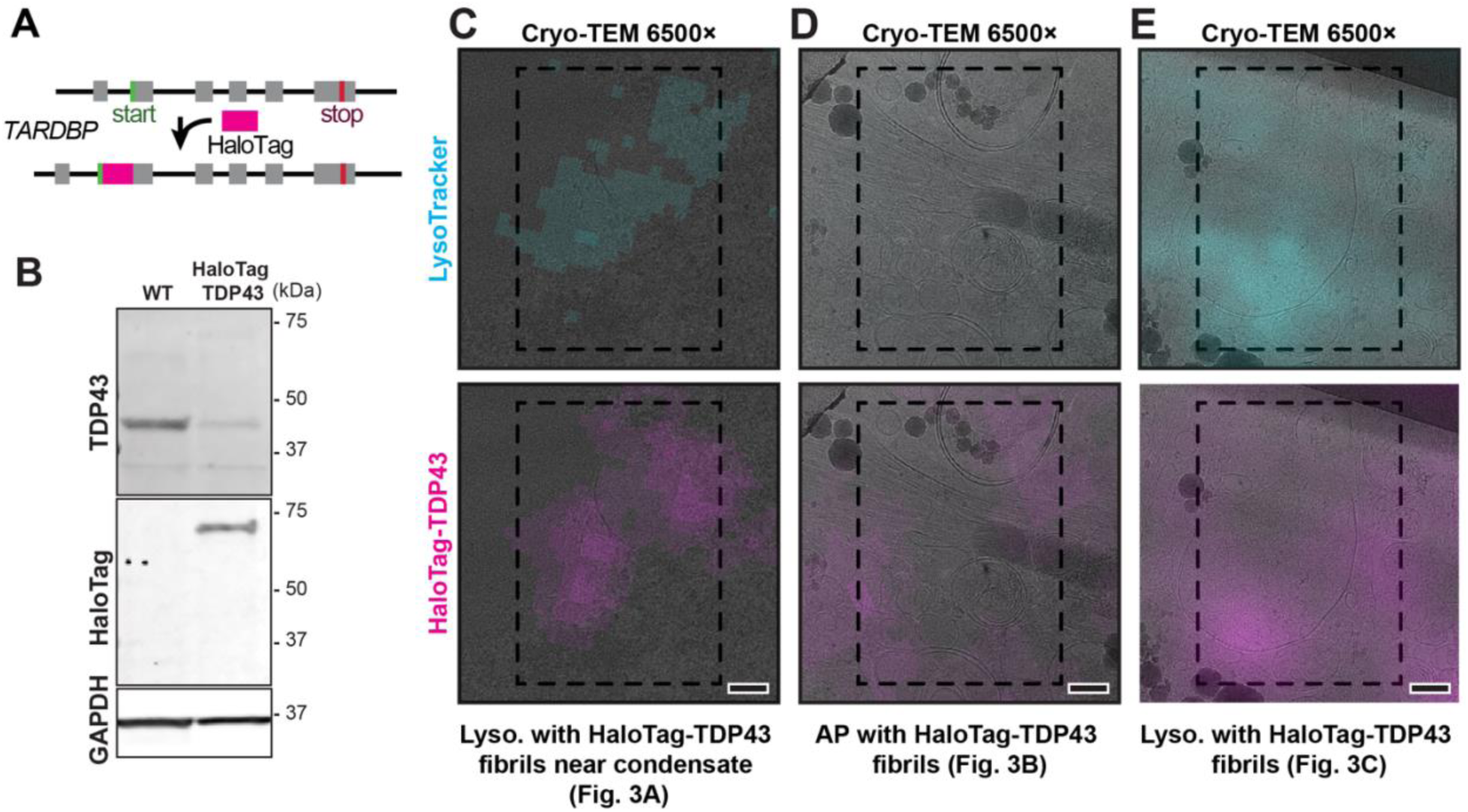
HaloTag-TDP43 iNeurons for *in situ* cryo-ET analysis. (**A**) Schematic for CRISPR/Cas9-based insertion of the HaloTag open reading frame immediately downstream of *TARDBP* start codon. (**B**) Immunoblots of unmodified (WT) and HaloTag-TDP43 iPSC lysates probed for TDP43, HaloTag, and GAPDH. (**C-E**) Fluorescence overlay of LysoTracker (cyan, top row) or HaloTag-TDP43 (magenta, bottom row) and the 6500× cryo-transmission electron microscopy (cryo-TEM) montage image of the lamella for tomograms shown in Figure 3A, Video S2 (C); Figure 3B, Video S3 (D); and Figure 3C, Video S4 (E). Tomography tilt series were collected in the areas of the cryo-lamella highlighted by the black dashed rectangle. Scale bar: 200nm

**Figure S5.**
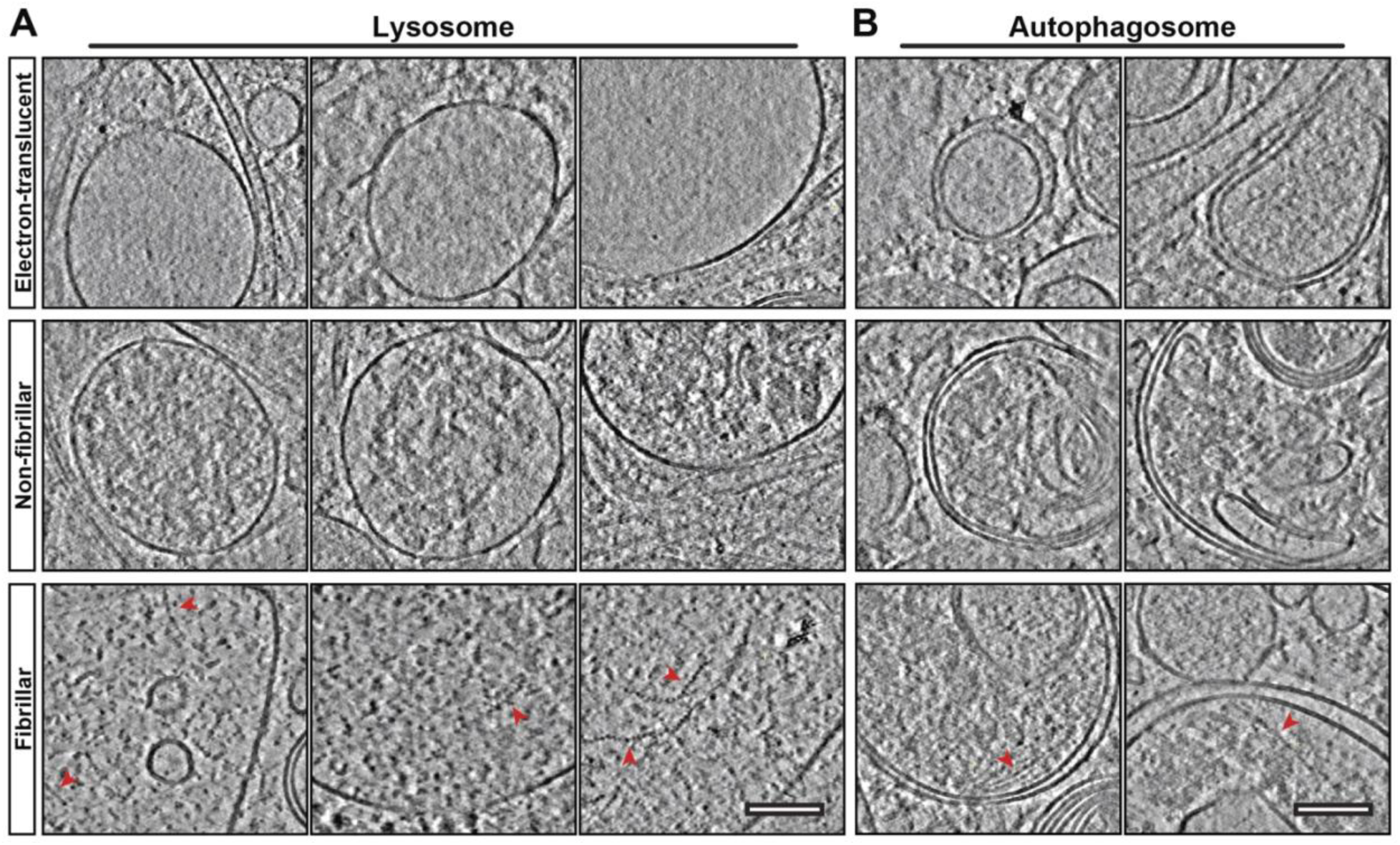
Gallery of lysosomes and autophagosomes from Halo-TDP43 iNeurons. (**A-B**) Representative slices through deconvolved tomograms obtained through cryo-CLEM workflow (Figure 2) of (GU)_6_ oligo treated Halo-TDP43 iNeurons showing electron-translucent (top row), non-fibrillar (middle row), and fibril-containing (bottom row) single-membrane (**A**) or double-membrane bound organelles (**B**). Red arrowheads highlight distinguishable TDP43 fibrils. Scale bar, 100nm.

**Figure S6.**
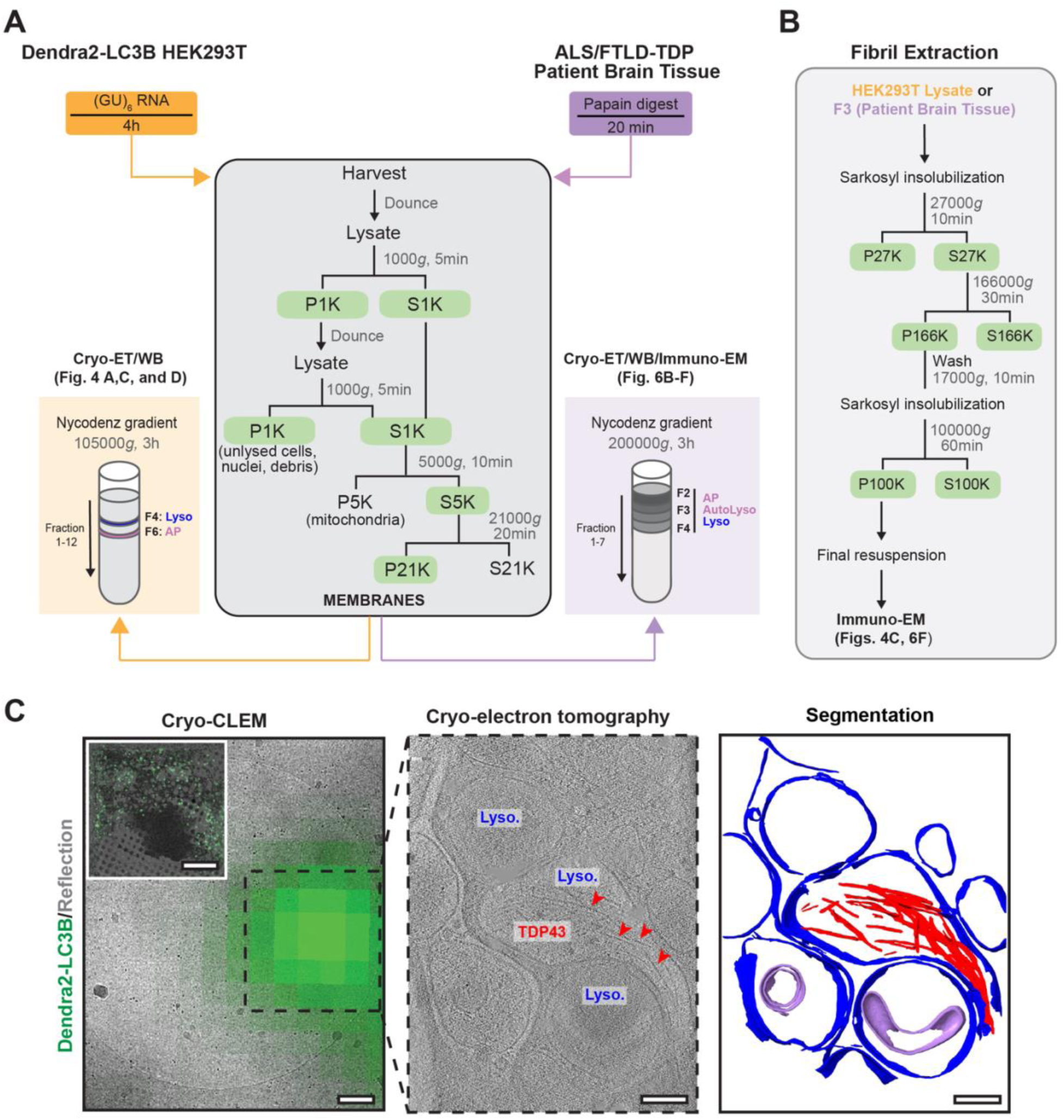
Isolation of autophagosomes and lysosomes from (GU)_6_-treated HEK293T cells or ALS/FTLD patient postmortem brain tissue. (**A**) Detailed schematic of gradient fractionation protocol for isolating autophagosomes (AP) and lysosomes (Lyso) from HEK293T cells (left) and ALS/FTLD patient postmortem brain tissue (right). (**B**) Schematic of fibril extraction protocol to isolate sarkosyl-insoluble TDP43 filaments from either HEK293T cells or ALS/FTLD patient postmortem brain tissue lysosomes. (**C)** Representative cryo-correlative light and electron microscopy (cryo-CLEM) images of autophagosome-enriched fraction 6 from Dendra2-LC3B HEK293T cells treated with (GU)_6_ for 4h. Overlay of the Dendra2-LC3B fluorescence signal with the reflection channel for a grid square (left, inset). Scale bar: 20μm. Overlay of Dendra2-LC3B fluorescence data with the 6500× cryo-TEM image of a hole within the grid square featured in the inset. Dashed box indicates the region of the hole with Dendra2-LC3B fluorescence signal targeted for tilt series acquisition (left). Scale bar: 200nm. Slice through a reconstructed tomogram showing a single membrane-bound Dendra2-LC3B positive lysosome containing TDP43 fibrils (middle). Scale bar: 200nm. Red arrowheads show the fibrils. The corresponding 3D segmentation (right) highlights lysosomes (blue), fibrils (red), and vesicles (lilac). Scale bar: 200nm.

**Figure S7.**
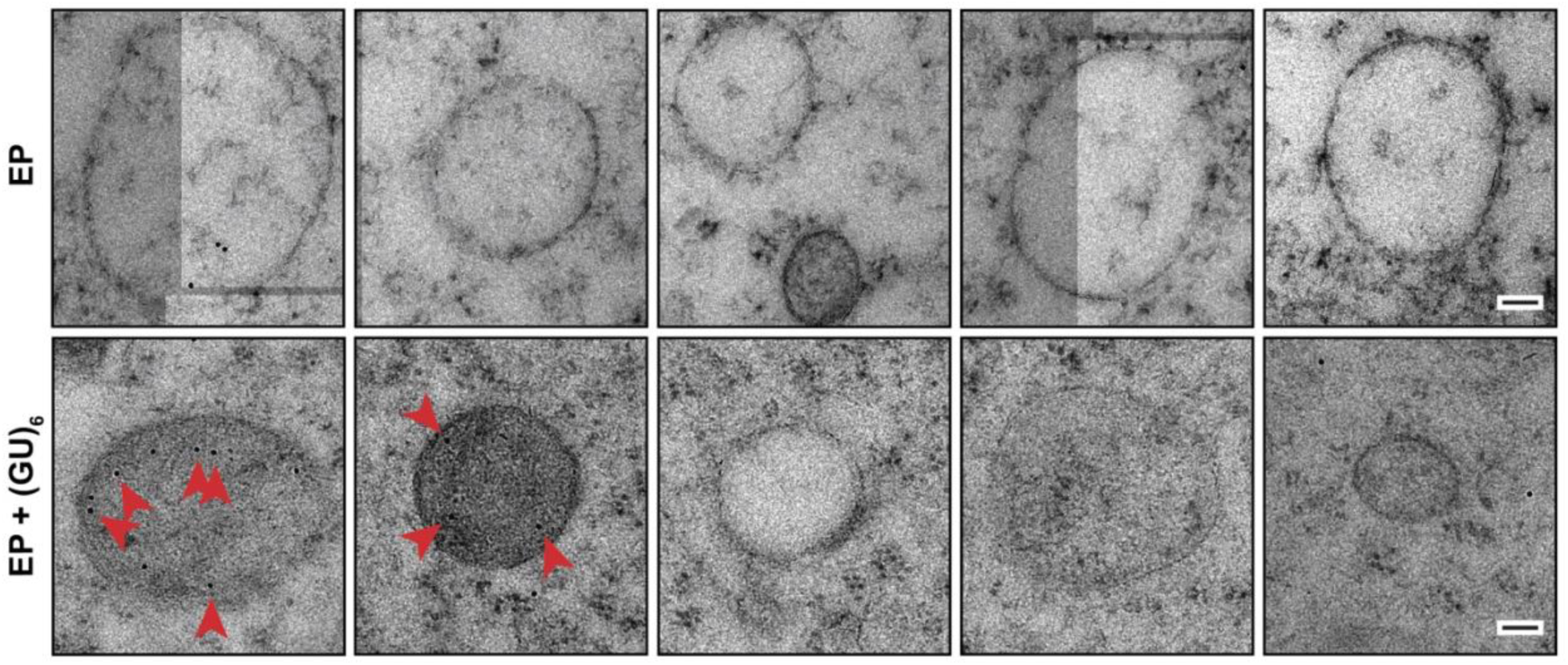
Immunogold labelling of HEK293T cell sections. (GU)_6_ oligonucleotide treatment leads to the accumulation of TDP43 in lysosomes in HEK293T cells. Representative transmission electron microscopy (TEM) images of single-membrane bound lysosomes from thin sections of Dendra2-LC3B HEK293T cells treated with EP (top) or EP and (GU)_6_ oligo (bottom) for 4h, showing 12nm gold beads. Colloidal gold beads conjugated to IgG (Goat anti-Rabbit) recognize the TDP43 antibody (10782-2-AP) and indicate the presence of TDP43. Scale bar: 100nm.

**Figure S8.**
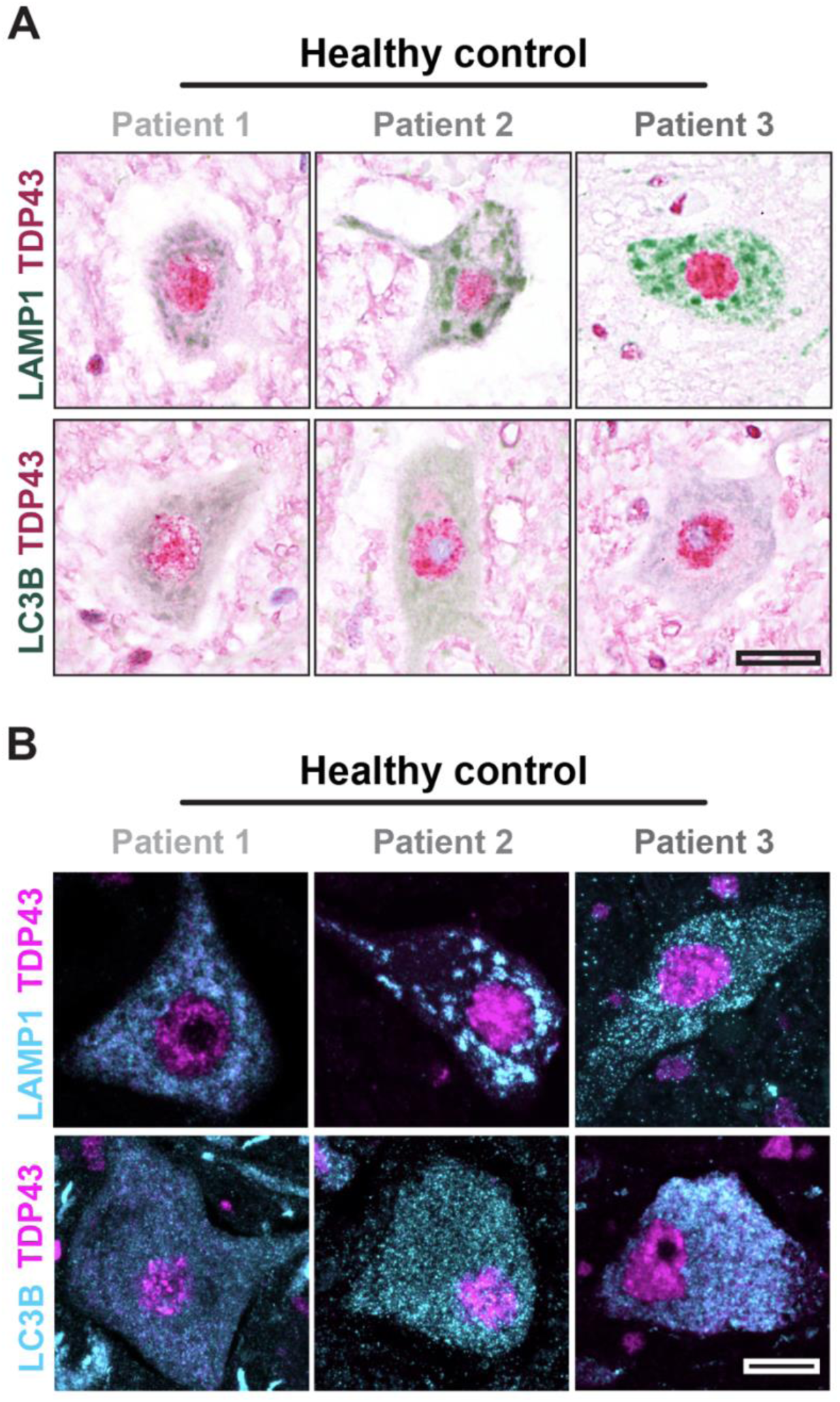
TDP43 and lysosome/autophagosome staining in control patient tissue. (**A**) Motor neurons from the spinal cords of multiple control patients (for simplicity, referred to as Patient 1, 2, and 3) underwent multiplexed immunohistochemistry for TDP43 (magenta), LAMP1 (green, top), and LC3B (green, bottom). Scale bar, 20µm. (**B**) Representative fluorescence micrographs of spinal neurons from 3 control patients immunostained for TDP43 (magenta), LAMP1 (cyan, top), or LC3B (cyan, bottom). Scale bars: 10 µm.

